# The mycobacterial ATP burst is a lysis artifact and serves as an assay for drug-induced cell wall damage

**DOI:** 10.1101/2025.10.24.684413

**Authors:** Claire V. Mulholland, Bei Shi Lee, Alexander Chong, Jinhua Cui, Kevin Pethe, Michael Berney

## Abstract

Antibiotics targeting the mycobacterial cell wall are a cornerstone of tuberculosis treatment. Understanding how these drugs facilitate bacterial killing has important implications for identifying novel therapeutic targets and optimizing drug combinations. Previous studies on mycobacteria using the BacTiter-Glo™ luminescence assay have reported increased ATP levels after treatment with cell wall inhibitors such as isoniazid, a phenomenon referred to as an ‘ATP burst’. This phenomenon has been proposed to contribute to drug-induced killing. In this study, we demonstrate that the ATP burst is not a biological response but rather an experimental artifact resulting from enhanced cell lysis induced by cell wall-targeting drugs. The addition of a bead-beating lysis step prior to luminescence measurement abolishes the ATP burst, enabling a more reliable assessment of intracellular ATP levels with this assay. Our findings thus establish a direct link between the ATP burst and the canonical mechanism of action of cell wall inhibitors. We further demonstrate the utility of this approach as a functional readout for identifying compounds that disrupt the mycobacterial cell wall and for screening synergistic or antagonistic interactions with known cell wall-targeting agents. Together, our results elucidate the mechanistic basis of the ATP burst and provide a valuable tool for antimycobacterial drug discovery and development.

## Introduction

*Mycobacterium tuberculosis* (*Mtb*) remains one of the world’s deadliest pathogens, causing more than one million deaths annually.^1^ In parallel, infections with non-tuberculous mycobacteria (NTM) have emerged as an escalating public health concern.^2^ In response to the urgent need for new therapeutic strategies against mycobacterial diseases, the past three decades have witnessed a marked expansion in molecular research aimed at elucidating the biology of these pathogens. A key aspect of this research has been the measurement of cellular ATP levels as a proxy for metabolic and bioenergetic state, for example, in studies of mycobacterial physiology and drug mechanisms of action,^3-7^ and high-throughput drug screening.^8-10^ Over the past two decades, luciferase-based assay kits, such as BacTiter-Glo™, have been widely adopted in mycobacterial research due to their affordability, rapid turnaround time, and ease of use. Notably, a search of PubMed Central for ‘BacTiter-Glo & Mycobacterium’ yields over 300 results, reflecting its widespread use. Recent studies using this assay have revealed an intriguing and unexpected phenomenon: treatment of different mycobacterial species (*Mtb, M. bovis* BCG, *M. abscessus*, and

*M. smegmatis*) with cell wall biosynthesis inhibitors leads to an apparent increase in ATP levels.^3,4,11-16^ This transient increase, described as an “ATP burst”, has been hypothesized to represent a lethal cellular process^4^ and to implicate electron transport chain perturbation in isoniazid-induced killing.^3^ However, it is still mechanistically unclear if this surge in ATP results from increased ATP production, decreased ATP hydrolysis, or an unknown mechanism, and how this might contribute to bacterial cell death.

Since ATP measurements based on the BacTiter-Glo™ method have become a standard in mycobacteriology, we sought to elucidate the mechanistic basis of the ATP burst phenomenon. Furthermore, the mycobacterial cell wall is a favorable target space for drug discovery,^17^ and cell wall inhibitors are central components of the multidrug treatments used to combat *Mtb* and NTM infections.^18,19^ Disruption of the mycobacterial cell wall can also improve the uptake and efficacy of other drugs.^20-25^ Conversely, drugs that target bioenergetics reduce the bactericidal efficacy of cell wall-targeting compounds.^3,4,14,16^ Thus, a better understanding of the mechanisms by which cell wall inhibitors lead to cell death is required to improve combination therapies and may reveal potential new drug targets.

In this work, we show that the ATP burst observed in the BacTiter-Glo™ assay is an experimental artifact resulting from synergistic lysis between cell wall-targeting drugs and the lysis buffer of the assay, leading to enhanced ATP release from *Mtb* and other mycobacteria. We demonstrate how this phenomenon can be exploited to identify compounds that target the cell wall and to detect potential synergistic and antagonistic interactions, providing a simple and rapid method to assess drug effects on the mycobacterial cell wall.

## Results

### The ATP burst is an experimental artifact caused by improved lysis following treatment with cell wall inhibitors

Mycobacteria have a thick and complex cell wall that is inherently resistant to lysis by conventional methods. The BacTiter-Glo™ assay relies on proprietary reagents to induce bacterial lysis; however, commercial systems are generally much less efficient at lysing mycobacteria compared to mechanical methods such as bead beating or heat inactivation.^26-28^ In our preliminary work, we detected no ATP burst in response to isoniazid when *Mtb* cells were heat-inactivated prior to the BacTiter-Glo™ assay (Fig. S1). Since heat inactivation leads to cell lysis, we hypothesized that the ATP burst might be related to lysis rather than an actual increase in cellular ATP levels. To test this systematically, we treated *Mtb* mc^2^6230^29^ with the cell wall inhibitors isoniazid and ethambutol, as well as the ATP synthesis inhibitor bedaquiline. ATP was then measured with the BacTiter-Glo™ assay using three different lysis methods: (1) ‘whole-cell’: incubating whole-cell culture with the BacTiter-Glo™ reagent for kit-driven lysis, (2) ‘beat’: bead beating whole-cell culture followed by a brief heat denaturation, or (3) ‘heat’: boiling whole-cell culture for 30 minutes at 100 °C. The lysate from both the heat and beat methods was spun down and then used in the BacTiter-Glo™ assay in the same manner as the whole-cell culture. Cells at high density (OD_600_ 0.33; ∼1 × 10^8^ CFU/ml) were treated at 50 × the minimum inhibitory concentration (MIC) to elicit robust responses and provide sufficient biomass for mass spectrometry analysis.

In agreement with previous reports,^3,4,11,13,14^ we observed a significant increase in the ATP signal for isoniazid and ethambutol treatment in the whole-cell assay (14- and 6-fold increase, respectively) (Fig. 1A). However, when cells were lysed by bead beating, no increase in ATP was observed, and instead, ATP levels were slightly lower compared to the untreated control (Fig. 1B). Concurrently, heat-inactivated samples also failed to show an ATP burst (Fig. S2B). Importantly, bead-beating of untreated cells increased the ATP signal by approximately 30-fold compared with the whole-cell assay (Fig. 1A–B), indicating ineffective lysis of mycobacterial cells by the BacTiter-Glo™ kit. We also tested whether isoniazid or ethambutol alone induced ATP release; however, neither drug substantially increased supernatant ATP (the signal was ≤2% of that observed in the whole-cell assay) (Fig. S2E). This indicates that the ATP burst is not a direct consequence of drug-induced lysis, but rather results from cell wall weakening, which enables effective lysis upon exposure to the BacTiter-Glo™ lysis buffer.

**Figure 1.**
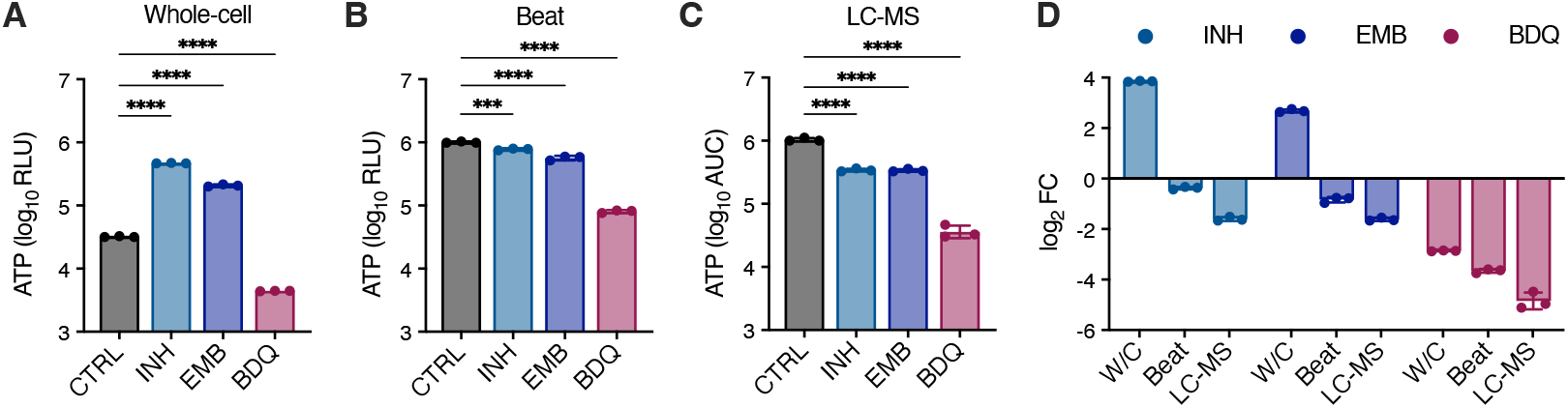
The ATP burst is an experimental artifact. *Mtb* mc^2^6230 at OD_600_ 0.33 was treated with isoniazid (INH, 3 μg/ml), ethambutol (EMB, 50 μg/ml), or bedaquiline (BDQ, 12.5 μg/ml) alongside untreated controls for 24 h (∼50 × MIC). **A–B.** ATP was measured using the BacTiter-Glo™ assay from either (A) whole-cell broth (‘W/C’) or (B) cells lysed by bead beating (‘Beat’). **C**. Intracellular ATP levels in the same cultures measured by LC-MS. **D**. Fold-change in ATP levels compared to untreated controls for the different methods shown in (A–C). Mean ± SD, *n* = 3 replicate cultures. \*\*\**P* < 0.001, \*\*\*\**P* < 0.0001; one-way ANOVA with Šídák’s multiple comparison test (MCT). See also Fig. S2.

We next sought to confirm relative ATP levels using an independent method and turned to liquid chromatography-mass spectrometry (LC-MS). ATP was extracted into water/acetonitrile/methanol buffer and measured on an Agilent 6545 Q-TOF mass spectrometer. No ATP burst was detected by this method (Figs. 1C and S3). Instead, ATP levels were significantly lower in isoniazid- and ethambutol-treated cultures compared to the untreated control. As expected, all methods detected significantly reduced ATP in cells treated with the ATP synthase inhibitor bedaquiline (Figs. 1 and S2).

To test if our findings also apply to other mycobacteria, we treated *M. abscessus* and *M. smegmatis*, two non-tuberculous mycobacteria, as well as the Gram-negative bacterium *Escherichia coli*, with the peptidoglycan biosynthesis inhibitor vancomycin at approximately 4–6 × their respective MICs,^30-32^ and measured ATP using the BacTiter-Glo™ whole-cell and beat assays. In agreement with our findings in *Mtb*, vancomycin treatment of *M. smegmatis* and *M. abscessus* significantly increased ATP in the whole-cell assay but not when cells were lysed by bead beating, and furthermore, bead beating increased the ATP signal in untreated cells by approximately 10- to 180-fold compared to the whole cell assay (Fig. 2A–C). Conversely, *E. coli*, which is efficiently lysed by the BacTiter-Glo™ lysis buffer, showed a decrease in ATP signal with vancomycin treatment in both assays, and the ATP signal was not increased by bead-beating (Fig. 2D).

**Figure 2.**
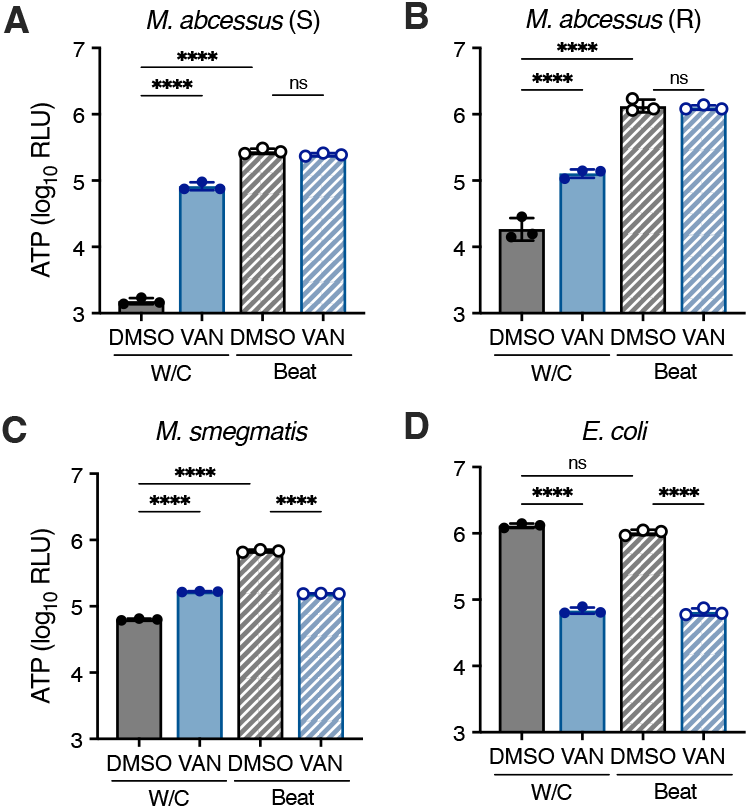
ATP release in the BacTiter-Glo™ whole-cell assay is impaired in mycobacteria due to the complex cell wall. **A–B.** *M. abscessus* smooth (S) and rough (R) morphotypes, and **C**. *M. smegmatis*, were treated for 4 h with 250 μg/ml vancomycin. **D**. *E. coli* was treated for 2 h with 500 μg/ml vancomycin. ATP was measured using the BacTiter-Glo™ whole-cell and beat assays. All cultures were treated at OD_600_ 0.033 alongside DMSO-only controls. Mean ± SD, *n* = 3 replicate cultures. \*\*\*\**P* < 0.0001; one-way ANOVA with Šídák’s MCT. These experiments were repeated with similar results.

Collectively, these findings demonstrate that the ATP burst observed in the BacTiter-Glo™ whole-cell assay is an experimental artifact. In the absence of prior weakening of the cell wall by drugs, the BacTiter-Glo™ lysis buffer alone is insufficient to effectively lyse mycobacterial cells. This has important implications for all applications of this assay in mycobacteria, as cell wall integrity and permeability can vary depending on growth conditions and phase,^33^ making assay results highly dependent on cell wall status. To overcome this limitation and minimize bias introduced by incomplete lysis, mechanical disruption by bead beating provides a more reliable method when using this assay.

### Whole-cell ATP screening identifies cell wall synthesis inhibitors

Having established the mechanistic basis of the ATP burst, we hypothesized that this response could serve as a functional assay to identify compounds that target the mycobacterial cell wall. In a proof-of-principle experiment, we screened a panel of 28 compounds with diverse mechanisms of action to confirm the specificity of the ATP burst to cell wall-targeting drugs and to adapt the assay for high-throughput. These included inhibitors of DNA replication, transcription, translation, folate biosynthesis, cell wall biosynthesis, and bioenergetics (Table 1). *Mtb* mc^2^6230 was treated with drugs at 10 μM and ATP was measured using the BacTiter-Glo™ whole-cell assay. After 24 hours, a strong ATP signal increase (>4-fold) specific to cell wall-targeting drugs was observed (Figs. 3A–B and S4), consistent with previous reports.^4^ Bioenergetics-targeting drugs showed their strongest effect at 48 hours (Fig. S4). The only cell wall drug not producing an ATP burst at 10 μM was D-cycloserine; however, at higher concentrations, an ATP burst was observed (Fig. 3D). We also observed a significant ATP increase with the serine esterase inhibitor tetrahydrolipstatin (THL) (Fig. 3A–B). THL has been reported to target lipid esterases involved in cell wall biosynthesis such as Pks13, TesA, and Ag85C,^34,35^ and has been shown to reduce levels of the cell wall lipids PDIM, SL-1, and TDM.^22,36^ However, unlike isoniazid and D-cycloserine, THL did not produce a strong sigmoidal dose-response curve (Fig. 3D), consistent with its broad nonspecific activity.^34,35^

**Table 1.**
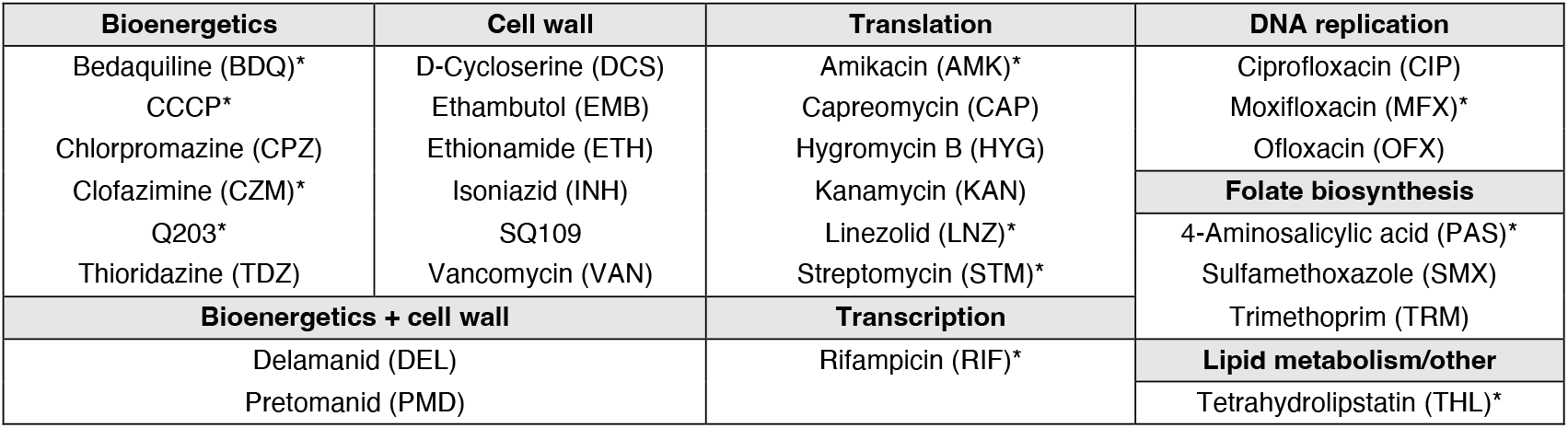
Reference compounds included in this study. All 28 compounds were included in the whole-cell ATP screen (Fig. 3A–B), and the 11 compounds indicated with an asterisk (*) were included in the cotreatment screen with isoniazid (Fig. 5A).

**Figure 3.**
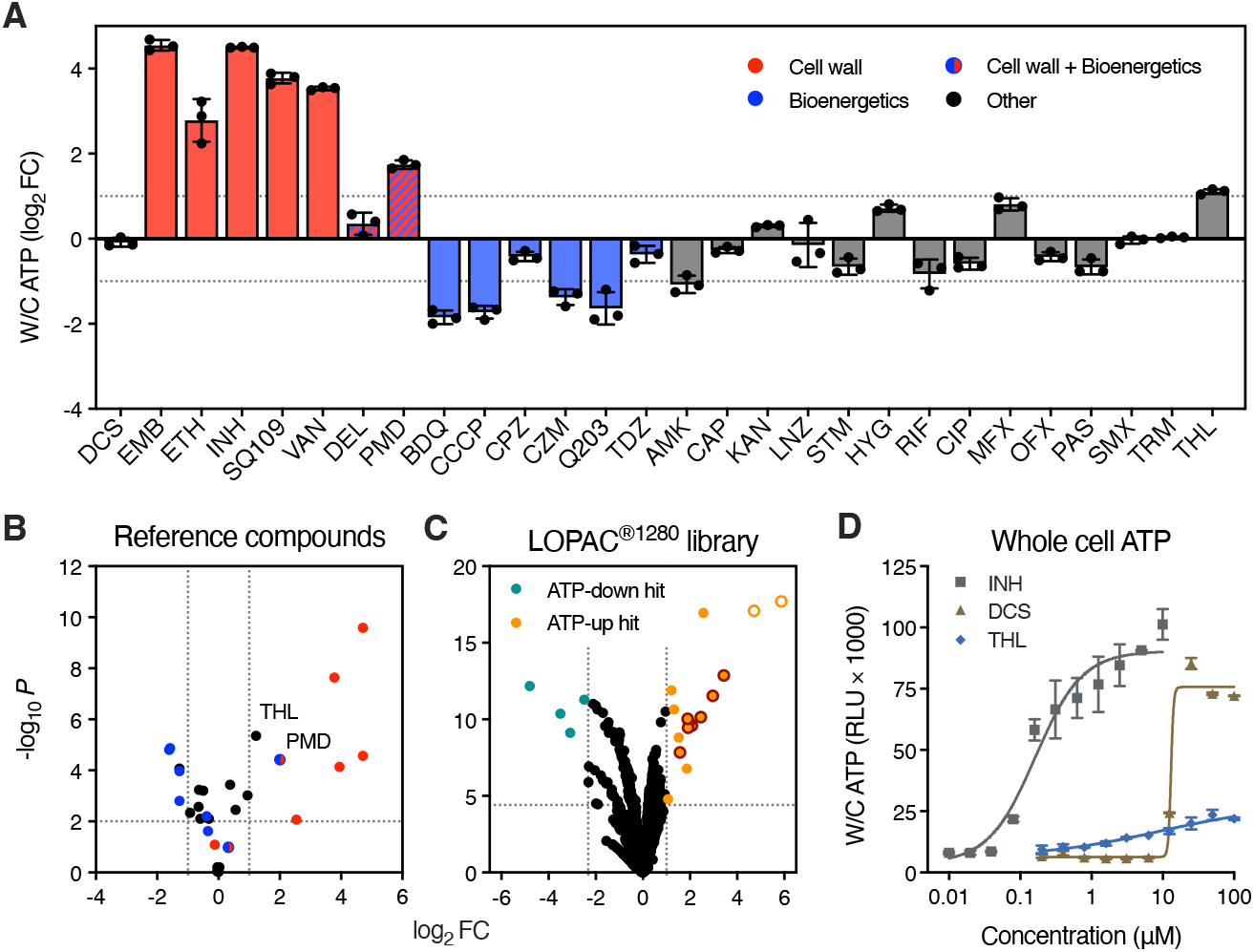
High-throughput ATP screening identifies compounds that target the *Mtb* cell wall. **A–B.** *Mtb* mc^2^6230 at OD 0.033 was treated for 24 h with 28 reference compounds (Table 1) at 10 μM alongside DMSO-only controls, and ATP was measured using the BacTiter-Glo™ whole-cell assay. (A) Log_2_ fold-changes relative to DMSO-only. Mean ± SD, *n* = 3 replicate wells. (B) Volcano plot for the experiment in (A). The dotted lines indicate a 2**-**fold change and a *P*-value of 0.01. See also Fig. S4. **C**. Volcano plot showing responses relative to DMSO-only controls for the Sigma LOPAC^®1280^ compound library screened using the same experimental setup in A–B. The dotted lines indicate a 2**-**fold change for “ATP-up” hits, an 80% decrease for “ATP-down” hits, and a *P*-value of 4 × 10^-5^ (Bonferroni cutoff, α = 0.05). Orange circles with dark red borders denote known cell wall inhibitors, and hollow orange circles denote compounds structurally related to ATP. See also Table 2. **D**. Dose-response curves for isoniazid (INH), D-cycloserine (DCS), and tetrahydolipstatin (THL). *Mtb* mc^2^6230 at OD 0.033 was treated for 24 h with increasing concentrations of each drug, and ATP was measured using the BacTiter-Glo™ whole-cell assay. Mean ± SD, *n* = 3 replicate wells. This experiment was repeated with similar results.

We then tested the feasibility of this approach using the Sigma LOPAC^®1280^ library of 1280 pharmacologically active compounds. This screen identified 16 compounds that induced a >2-fold increase in the ATP signal (“ATP-up” hits) (Fig. 3C, Table 2). Seven of these hits were cephalosporins, which are β-lactam antimicrobials that inhibit peptidoglycan biosynthesis, validating our approach. Furthermore, the hit compound nialamide is a derivative of isoniazid and accordingly showed cross-resistance in an isoniazid-resistant mutant (Fig. S5). Two compounds: P1,P4-di(adenosine-5’)tetraphosphate ammonium and 2-chloroadenosine triphosphate tetrasodium, were structurally related to ATP and increased the BacTiter-Glo™ signal independent of *Mtb* and were therefore excluded from further analyses. This left six novel ATP-up hits not previously known to target the cell wall. We assessed the ability of these compounds to inhibit *Mtb* growth, along with nialamide, the cephalosporin cefotaxime, and four compounds that were found to lower ATP by >80% (“ATP-down” hits) (Fig. 3C, Table 2). Three of the six novel ATP-up hits – Bay 11-7082, Bay 11-7085, and L-687,384, as well as nialamide, cefotaxime, and all four ATP-down hits, inhibited the growth of both *Mtb* mc^2^6230 (Fig. 4A–B) and virulent *Mtb* H37Rv (Fig. S6) with MIC_90_ concentrations between 0.8–50 μM, demonstrating the ability of this approach to identify compounds with antitubercular activity in virulent and avirulent strains of *Mtb*. Furthermore, six of these compounds: nialamide, L-687,384, and all four ATP-down hits, have previously been shown to inhibit *M. bovis* BCG^37^ (Table 2). Antibacterial compounds in the library with other known mechanisms, such as doxycycline and ofloxacin, which target protein synthesis and DNA gyrase, respectively, did not come up as hits in our screen, highlighting the specificity of the setup to identify cell wall-targeting compounds.

**Table 2.**
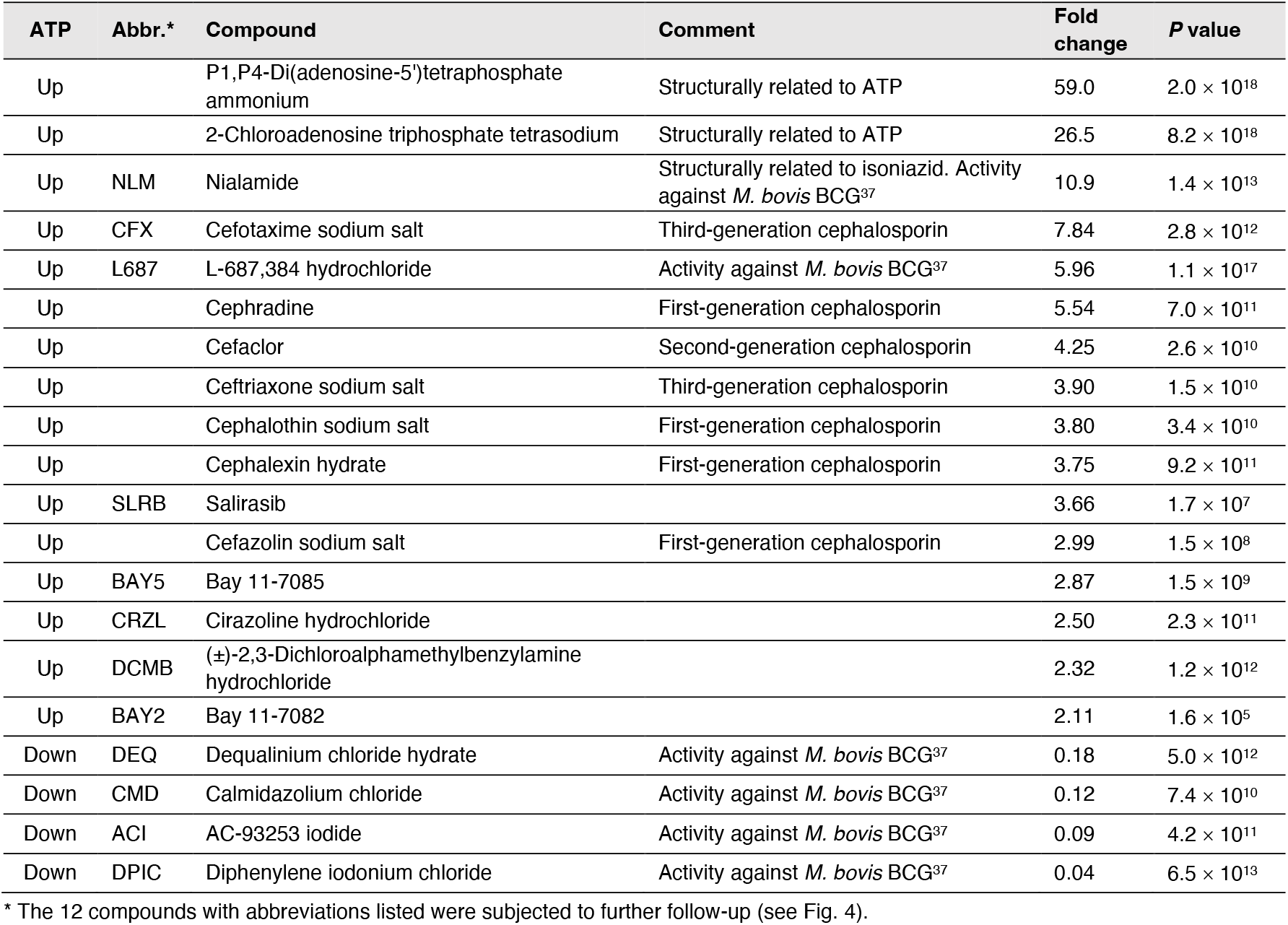
Sigma LOPAC^®1280^ compound library whole-cell ATP screening hits. Small molecule hits identified in the BacTiter-Glo™ whole-cell assay that either significantly increased the ATP signal by >2-fold (“ATP-up” hits) or significantly decreased the ATP signal by >80% (“ATP-down” hits). See also Fig. 3C.

**Figure 4.**
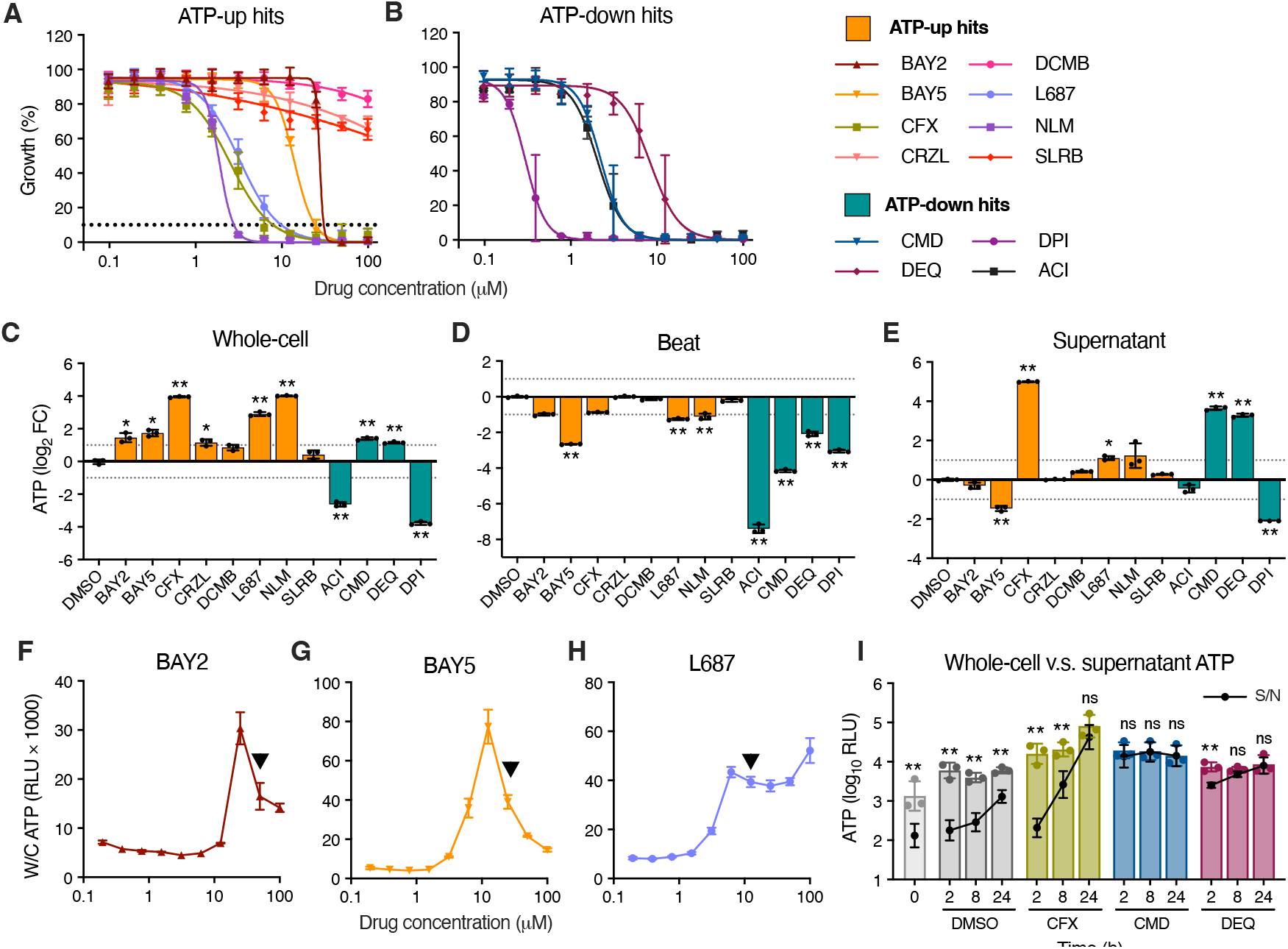
Validation of ATP screening hits from the LOPAC compound library. **A–B.** MIC assays for *Mtb* mc^2^6230 with selected LOPAC library hits (see Table 2). Growth was measured by OD_600_ and normalized to no-drug control wells. Mean ± SD, *n* = 4 replicate wells from two independent experiments. See also Fig. S6. **C–E**. Follow-up of ATP responses to LOPAC hits. Cultures at OD_600_ 0.033 were treated for 24 h at 20 μM, and ATP was measured using (C) the BacTiter-Glo™ whole-cell assay, (D) the bead-beating assay, and (E) in the culture supernatant. Mean ± SD, *n* = 3 replicate cultures. \**P* < 0.05, \*\**P* < 0.001 and ≥2**-**fold change; unpaired *t*-test with Holm-Šídák’s MCT. **F–H**. ATP whole-cell assay dose-response curves for (F) Bay 11-7082, (G) Bay 11-7085, and (H) L-687,384. Black arrows indicate the MIC_90_. Mean ± SD, *n* = 3 replicate wells. **I**. Time course of ATP responses in the BacTiter-Glo™ whole-cell assay (bars) and in the culture supernatant (S/N, black circles and line) following treatment with 20 μM cefotaxime (CFX), calmidazolium (CMD), or dequalinium (DEQ). Mean ± SD, *n* = 3 means from replicate experiments performed in either duplicate or triplicate. \*\**P* < 0.001 and ≥2**-**fold change between supernatant and whole-cell ATP signal; unpaired *t*-test with Holm-Šídák’s MCT. For experiments in (C– H), cultures at OD_600_ 0.033 were treated for 24 h or for the duration indicated in (I). All experiments were repeated with similar results.

We next followed up on these compounds by measuring ATP levels using our bead-beating assay to confirm that they indeed weaken the cell wall rather than increase ATP through an unrelated mechanism. The three novel ATP-up compounds that inhibited *Mtb* growth showed a >2-fold ATP increase in the whole-cell assay, which was accompanied by an ATP decrease in the beat assay, consistent with impaired cell wall integrity and increased ATP release (Fig. 4C–D). The three compounds that failed to inhibit growth showed a weak or no response in the whole-cell assay follow-up and mirrored DMSO in the beat assay, consistent with the absence of antimycobacterial activity. Interestingly, unlike classical cell wall inhibitors, which produced strong sigmoidal dose-response curves (Fig. 3D), BAY2 and BAY5 exhibited dose-response peaks at 0.5 × MIC_90_, suggesting these compounds might have multiple modes of action (Fig. 4F– G). We note that these two compounds, together with those lacking antimycobacterial activity and several cephalosporins, showed weaker responses in the initial screen (2–4-fold changes) (Table 2). This suggests that a 2-fold detection threshold may sit near the cutoff between meaningful activity and noise, while still detecting compounds with weak cell wall-targeting effects or low potency.

Surprisingly, in our follow-up assay, two of the ATP-lowering hits – calmidazolium (CMD) and dequalinium (DEQ) increased the ATP signal in the whole-cell assay, though it was substantially lowered in the beat assay (Fig. 4C–D). Both compounds also rapidly increased supernatant ATP levels, indicating cell lysis (Fig. 4E,I). In agreement with our results, dequalinium is reported to induce a range of effects on microbial cells, including inhibition of the F_1_-ATPase,^38^ and has been suggested to affect cell membrane permeability, potentially leading to lysis.^39^ Cefotaxime also substantially increased supernatant ATP after 24 hours, though not at earlier time points (Fig. 4E,I). These results demonstrate the application of the BacTiter-Glo™ assay for high-throughput identification of cell wall inhibitors as well as the versatility of this assay to probe the effects of antitubercular compounds on the *Mtb* cell wall and bioenergetics.

### Detection of antagonistic and synergistic interactions with cell wall drugs

Compounds that disrupt cellular bioenergetics result in reduced ATP production.^40^ Inhibitors of energy metabolism have been shown to antagonize cell wall-targeting compounds, suppressing the ATP burst and reducing killing,^3,4,14,16^ leading some authors to conclude that excessive ATP contributes to the lethality of cell wall drugs.^4^ Consistent with these studies, we observed suppression of the isoniazid-induced ATP increase in the whole-cell assay and reduced killing following co-treatment with the bioenergetics inhibitors bedaquiline, Q203, and CCCP (Fig. S7A–B). Conversely, the BacTiter-Glo™ beat assay showed decreased ATP in cultures treated with isoniazid alone, and this was further depressed by cotreatment with bioenergetics inhibitors (Fig. 6C). Considering that the ATP burst reflects disruption of the mycobacterial cell wall (Figs. 1 and 2), these data indicate that cotreatment with bioenergetics inhibitors leads to a reduction in the canonical activity of cell wall inhibitors rather than mitigating “death by ATP”.

We speculated that this antagonistic interaction could be partly related to general metabolic slowdown, in which case it may not be specific to bioenergetics inhibitors. To test this, we cotreated *Mtb* mc^2^6230 with isoniazid and 11 compounds with varying modes of action (Tables 1 and S1). To improve experimental throughput, we used 96-deepwell blocks incubated with vigorous shaking, which supports much faster *Mtb* replication rates than conventional 96-well microtitre plates (Fig. S8). Cotreatment with bioenergetics inhibitors (bedaquiline, Q203, CCCP, and clofazimine), as well as the protein synthesis inhibitor linezolid, substantially reduced the ATP signal at both doses tested compared to isoniazid alone, while THL significantly increased it (Figs. 5A), pointing to potential antagonistic and synergistic effects, respectively. We followed up on these interactions by comparing ATP levels in the BacTiter-Glo™ whole-cell and beat assays and plated for viability. Linezolid cotreatment significantly decreased isoniazid-induced ATP release and reduced killing by ∼1-log, consistent with an antagonistic effect (Fig. 5B–D). This interaction was not detected in the checkerboard assay (FICI = 1.0, Table S2); consequently, this effect would be missed if only the checkerboard assay was performed. Unlike bioenergetics inhibitors which lowered ATP in the beat assay (Fig. S7C), linezolid alone resulted in a small but significant increase in ATP (1.4-fold, *P* = 0.0002), and cotreatment did not further lower ATP compared to isoniazid alone (Fig. 5C). Given that protein biosynthesis is one of the most energetically costly cellular processes,^41^ this increase may result from an immediate reduction in ATP consumption upon translation arrest. In contrast, THL cotreatment significantly increased ATP in the whole-cell assay compared to isoniazid alone, with no difference in the beat assay, indicating greater ATP release in the cotreatment; however, this did not translate to an effect on killing (Fig. 5B–D). We again saw a small but significant increase in ATP in the whole-cell assay for THL alone (Figs. 3 and 5B), suggesting this may be an additive effect, consistent with what we observed in the checkerboard assay (FICI = 0.71, Table S2). Furthermore, isoniazid alone kills *Mtb* very rapidly,^42^ therefore, further improvement of isoniazid-induced killing might not be possible at the dose used.

**Figure 5.**
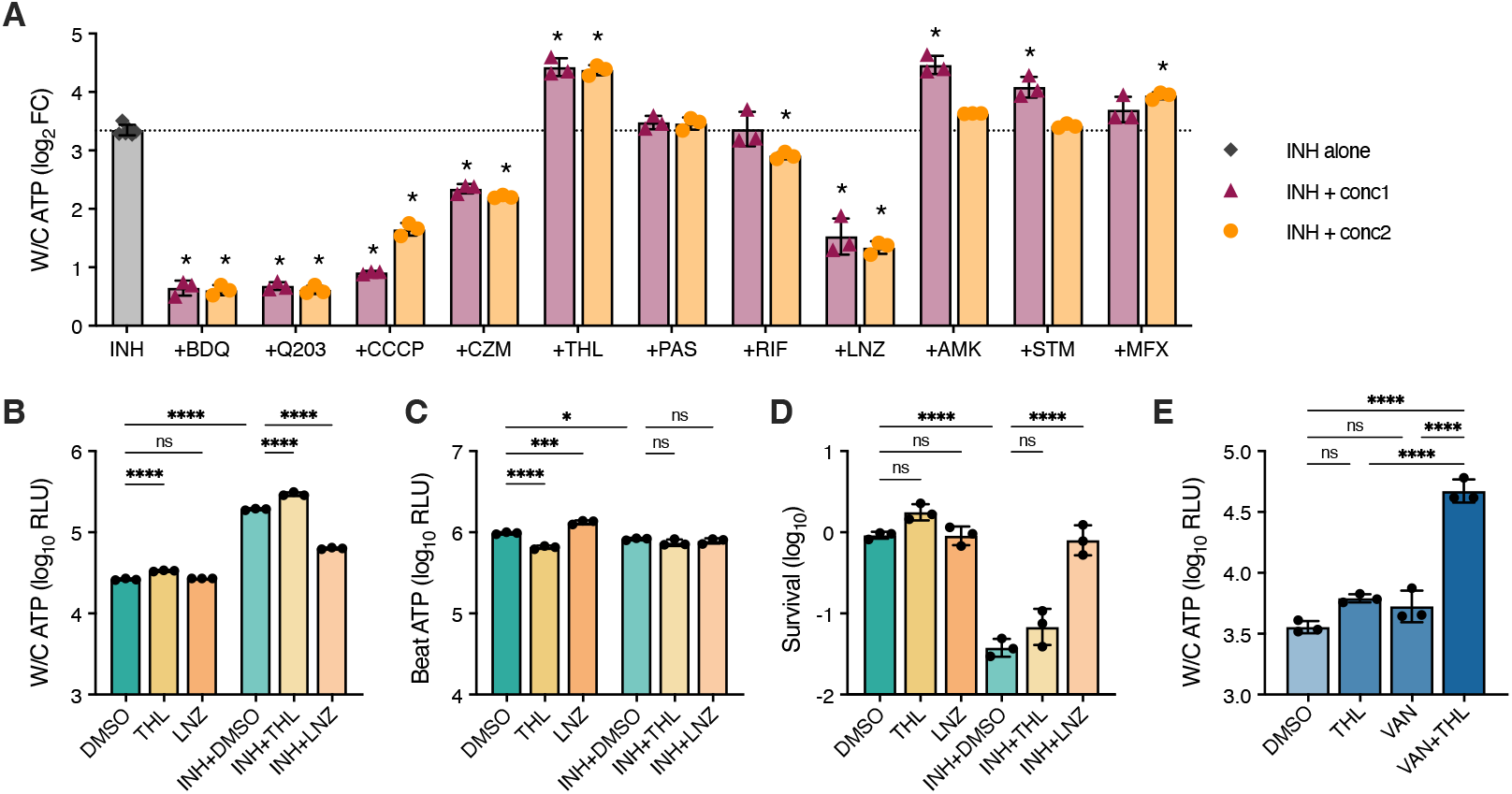
ATP screening identifies antagonistic and synergistic interactions with cell wall inhibitors. **A**. *Mtb* mc^2^6230 at OD 0.33 was treated for 24 h with isoniazid alone (INH, 0.1 μg/ml) or in combination with the indicated compounds at 2–4 × and 5–10 × MIC_90_ (“conc1” and “conc2,” respectively) (see Tables 1 and S1). Cultures were incubated in 96-deepwell blocks with fast shaking to maximize growth (see Fig. S8). ATP was measured using the BacTiter-Glo™ whole-cell assay. Data are shown as log_2_ fold-changes compared to the DMSO-only control. Mean ± SD, *n* = 6 replicate wells for DMSO and isoniazid, and *n* = 3 for cotreatments. \**P* < 0.001 vs. isoniazid alone; two-way ANOVA with Dunnett’s MCT. This experiment was performed once. **B–D**. Follow-up of putative antagonistic and synergistic effects identified in (A). Cultures at OD 0.33 were treated for 24 h with isoniazid (INH, 0.1 μg/ml), linezolid (LNZ, 2 μg/ml), and tetrahydrolipstatin (THL, 50 μg/ml) as indicated, and ATP was measured using (B) the BacTiter-Glo™ whole-cell and (C) beat assays, and (D) plated for CFU to assess viability. The starting cell density was 3 × 10^8^ CFU/ml. Mean ± SD, *n* = 3 replicate cultures. \**P* < 0.05, \*\*\**P* < 0.001, \*\*\*\**P* < 0.0001; one-way ANOVA with Šídák’s MCT. **E**. Cotreatment with tetrahydrolipstatin and vancomycin (VAN). PDIM(+) *Mtb* mc^2^6230 at OD 0.033 was treated with 1 μg/ml of each drug alone and in combination for 24 h, and ATP was measured using the BacTiter-Glo™ whole-cell assay. Cultures were treated in 96-well microtiter plates and supplemented with 0.1 mM propionate to support PDIM production. See also Fig. S10. \*\*\*\**P* < 0.0001; one-way ANOVA with Šídák’s MCT. Mean ± SD, *n* = 3 replicate wells. The experiments in B–E were repeated with similar results.

Next, we determined if the status of the cell wall lipid phthiocerol dimycocerosate (PDIM) impacts our findings. Since beginning this work, we established that our *Mtb* mc^2^6230 stock is a predominantly PDIM-negative mixed population and have re-isolated a pure PDIM-positive clone.^25^ In PDIM-positive *Mtb* mc^2^6230, isoniazid induced an artifactual ATP burst in the BacTiter-Glo™ whole-cell assay, both in standard 7H9 and propionate-supplemented PDIM maintenance media^25^ (Fig. S9). These results verify the applicability of this approach to PDIM-positive strains.

Finally, as a proof of concept, we used this approach to detect a previously described synergy between THL and vancomycin.^22^ Vancomycin is a large antibiotic whose efficacy is significantly affected by cell wall permeability and PDIM levels.^21,24,25^ THL synergizes with vancomycin by lowering PDIM production and impairing cell wall integrity.^22^ As both compounds cause an ATP burst, we first determined concentrations of vancomycin and THL that produced minimal effects on the BacTiter-Glo™ whole-cell ATP signal in PDIM-positive *Mtb* (Fig. S10). We found that at 1 μg/ml, both drugs induced only a ∼1.5-fold increase compared to the DMSO control; however, when used in conjunction, the ATP signal increased 13-fold, denoting a strong potentiation of cell wall effects (Fig. 5E). This is consistent with previous work showing synergism between THL and vancomycin^22^ and demonstrates the potential of this approach to screen for such interactions.

## Discussion

One key distinguishing feature of mycobacteria is their complex cell wall. This contributes to their intrinsic resistance to many antibiotics, their ability to persist in the host, and makes them notoriously difficult to lyse.^33^ As such, commercial kits that work well on other species with less complex cell walls, such as gram-negative bacteria, are much less efficient against mycobacteria, which are better lysed using more aggressive mechanical methods, such as bead beating or heat inactivation.^26-28^ Accordingly, we discovered that the previously reported ATP burst phenomenon^3,4,11-16^ is an artifact of improved cell lysis facilitated by drugs that weaken the cell wall. This is in line with Koul *et al*. 2014, who did not observe an increase in ATP in *Mtb* after 24 hours of isoniazid treatment using heat inactivation and an ATP bioluminescence assay kit.^5^ While our findings highlight the need for mechanical lysis to improve the accuracy and reliability of BacTiter-Glo™-based ATP measurements in mycobacteria, they also reveal a valuable research application of the ATP burst phenomenon. Our data establish a direct link between the ATP burst and the canonical mechanism of action of cell wall inhibitors, providing a rapid functional readout that can be leveraged in several ways: (1) to assess whether compounds with unknown mechanisms affect the mycobacterial cell wall; (2) as a high-throughput screening tool to identify new cell wall inhibitors from large compound libraries; and (3) to evaluate potential synergistic or antagonistic interactions with known cell wall-targeting drugs. An increase in ATP relative to controls (typically >2–4-fold) can be interpreted as evidence of a weakened cell wall and subsequently validated using the complementary bead-beating assay to quantify ATP in fully lysed cultures. Additional analyses, such as measuring extracellular ATP and generating dose-response curves, can further support and refine conclusions regarding cell wall-targeting activity. In optimizing our screen, we found that while cell wall targeting effects were best captured after 24 hours, the strongest response for bioenergetics inhibitors was observed after 48 hours. To enhance the assay’s utility as a dual-purpose screen, an additional later time point could be included to better capture ATP-lowering effects. However, extended incubation may increase the potential for confounding factors, such as variability due to differences in replication rates and cell death.

Our bead-beating protocol presents an approach to use the BacTiter-Glo™ kit to more reliably assess ATP levels in mycobacterial cultures by removing the bias of incomplete lysis. This approach may also increase assay sensitivity when assessing bioenergetic inhibitors, as incomplete lysis reduces the effective dynamic range. However, for a more detailed assessment of cellular bioenergetics, it may be necessary to determine the ratio of ATP, ADP, and AMP, as this provides a better representation of functional energy homeostasis than ATP level alone.^43^ This can be done, for example, by mass spectrometry.^44^ Alternatively, in recent years, there have been several reports using genetically encoded biosensors to measure mycobacterial ATP abundance^45-47^ or the ATP/ADP ratio.^48,49^ This enables real-time ATP(/ADP) tracking at the single-cell level with improved spatiotemporal resolution and avoids normalization challenges related to differences in biomass and cell viability. While these assays reliably detect reduced ATP levels in response to bioenergetic inhibitors,^45,46,48^ their performance with isoniazid has been inconsistent, with studies reporting ATP depletion in *M. smegmatis*,^46^ increased ATP in *Mtb*,^45^ and no effect on the ATP/ADP ratio in *Mtb*.^48^ This might be due to the use of different sensors in these studies, different strains, varying assay conditions, or sensitivity issues. Further improvement and standardization of ATP biosensors will provide a valuable tool for studying bacterial bioenergetics.

Our findings argue that bioenergetics inhibitors reduce the on-target activity of isoniazid, fitting with a model whereby drug-induced ATP depletion renders cells more tolerant to isoniazid due to decreased target activity, rather than by inhibiting a toxic ATP surge. This is in line with previous work correlating lowered ATP levels with increased drug tolerance and persistence in *Mtb*.^47^ Conversely, the antagonistic effect observed with linezolid was not associated with reduced ATP. Together with our previous work showing that bacteriostatic effects alone are insufficient to protect against isoniazid-induced killing,^14^ these observations support the notion that suppression of isoniazid-mediated killing is more complex than simply blocking replication.^14^

Independent of the specific mechanisms involved, our work demonstrates how whole-cell ATP luminescence screening can be employed to detect novel synergistic and antagonistic drug interactions with cell wall inhibitors. We anticipate that this could be further expanded, for example, to screen for nutrient or genetic interactions. This approach also enabled us to detect interactions missed by the conventional checkerboard assay. A similar discordance between bactericidal activity and growth inhibition has been reported in *M. abscesscus* for bedaquiline and β-lactams, which also have an antagonistic interaction in the BacTiter-Glo™ whole-cell assay.^16^ The ability to screen for antagonistic interactions can be particularly valuable in early drug development, helping to identify promising combinations as well as those that should be avoided. This approach could also be used to improve the potency of cell wall drugs whose activity is limited, for example, due to degradation (e.g. β-lactams) or poor uptake (e.g. vancomycin). β-lactams are a largely untapped resource for mycobacterial treatment, primarily due to the presence of chromosomally encoded β-lactamases that confer intrinsic resistance.^50^ Furthermore, in addition to canonical D,D-transpeptidases, most peptidoglycan cross-links in mycobacteria are 3→3 linkages catalyzed by non-classical L,D-transpeptidases, which are generally not inhibited by these drugs.^51^ Approaches to circumvent these limitations include using β-lactamase inhibitors^52,53^ or combinations of different β-lactam subclasses to target both L,D- and D,D-transpeptidases.^54^ ATP screening could provide a simple, high-throughput method for identifying novel β-lactam combinations, potentially aiding the development of more effective tuberculosis and NTM treatment regimens.

Finally, the high-throughput cultivation method developed in this work, enabling *Mtb* to be grown in 96-well plates at growth rates comparable to inkwells or roller bottles, will be a valuable new tool for tuberculosis research in general.

In conclusion, our work provides a mechanistic explanation for the ATP burst detected with the BacTiter-Glo™ kit in the presence of cell wall-targeting compounds, directly linking this phenomenon to the canonical mode of action of these drugs. We show that the simplicity, speed, low cost, and high-throughput compatibility of this approach make it a valuable addition to the antimycobacterial drug discovery and development toolkit. Moreover, we propose practical solutions to more accurately determine cellular ATP levels in mycobacteria by addressing the confounding effects of incomplete cell lysis.

## Materials and Methods

### Bacterial strains, culture conditions, and reagents

*Mtb* mc^2^6230 (H37Rv Δ*RD1* Δ*panCD*)^29^ and *M. smegmatis* mc^2^155^55^ strains were obtained from laboratory stocks. A pure PDIM-positive mc^2^6230 clone (AE1601) was previously isolated from our mc^2^6230 stock.^25^ PDIM-positive *Mtb* H37Rv (AE1001) was isolated following C57BL/6 mouse passage using previously described methods.^25^ *M. abscesses* (ATCC 19977) and *E. coli* K-12 (ATCC 10798) were obtained directly from ATCC. *Mtb* strains were cultured in Middlebrook 7H9 broth supplemented with 10% (v/v) OADC (0.6 g/l sodium oleate, 50 g/l bovine serum albumin fraction V, 20 g/l dextrose, 40 mg/l catalase, 8.5 g/l sodium chloride), 0.2% (v/v) glycerol, and 0.05% (v/v) tyloxapol, with the addition of 24 mg/l D-calcium pantothenate for *Mtb* mc^2^6230. PDIM-positive strains were additionally supplemented with 0.1 mM sodium propionate to maintain and enhance PDIM production.^25^ *M. abscesses* and *M. smegmatis* were cultured in Middlebrook 7H9 broth supplemented with 10% (v/v) Middlebrook ADC enrichment, 0.2% (v/v) glycerol, and 0.05% (v/v) Tween 80. Middlebrook 7H10 agar supplemented with 10% (v/v) OADC and 0.5% (v/v) glycerol was used as a solid medium for plating mycobacterial strains, with the addition of 24 mg/l D-calcium pantothenate for *Mtb* mc^2^6230. *E. coli* K-12 was cultured in LB broth, and LB agar was used for plating. Supplier information for media components and supplements are listed in Table S3. For *Mtb* experiments, fresh starter cultures were inoculated from frozen seed stocks, and for *M. abscesses, M. smegmatis*, and *E. coli*, single colonies were picked from agar plates. All strains were subcultured once, grown to logarithmic phase (OD_600_ ∼0.8), and then diluted to the required density for each experiment. Mycobacterial cultures were grown at 37 ºC with gentle shaking at 100 rpm for BSL2 strains and 80 rpm for BSL3. *E. coli* was incubated at 37 ºC with shaking at 200 rpm.

### BacTiter™ Glo ATP assays

The BacTiter-Glo™ Microbial Cell Viability Assay (Promega) was used to quantify ATP following treatment with antimicrobial compounds (see Table S4). Total DMSO concentration was kept ≤1% for all experiments. For whole-cell ATP assays, 25 μl of bacterial culture and 25 μl of the BacTiter-Glo™ reagent were combined in 96-well white opaque plates. Plates were incubated in the plate reader in the dark with shaking for 5 min, and then luminescence was measured using either a FLUOstar Omega or VANTAstar Microplate Reader (BMG Labtech). The LOPAC screen was measured using a Cytation-3 Cell Imaging Multiple-Mode Reader (LabX) with a 1 sec integration to capture luminescence. To pre-lyse cells by bead beating (‘beat’ assay), 250 μl of bacterial culture was added to approximately 250 μl of 0.1 mm zirconia/silica beads (BioSpec) and then lysed using a Precellys Cryolys Evolution bead beater (Bertin Technologies) cooled to 0 °C for three 20 sec cycles at 6,800 rpm with a 30 sec pause between cycles. Lysed samples were boiled at 100 °C for 5 min to denature cellular enzymes, then briefly cooled on ice before centrifugation at 4 °C for 5 min at 13,000 rpm. Heat lysis (‘heat’ assay) was performed by boiling 250 μl culture for 30 min at 100 °C and then pelleting as above. 25 μl of the lysate from either the beat or heat extractions and 25 μl of the BacTiter-Glo™ reagent were added to white 96-well plates, which were incubated and measured as for whole-cell samples. To quantify supernatant ATP, 250 μl of bacterial culture was centrifuged for 3 min at 13,000 rpm. The supernatant was then passed through a 0.22 μm filter, and ATP in the filtrate was measured as for whole-cell culture and lysates. Calibration curves of ATP chemical standards (Sigma) prepared in water were used to determine the linear range.

### ATP quantification by LC-MS

*Mtb* mc^2^6230 cultures at OD600 0.33 were treated with antimicrobial compounds as specified for 24 h in 10 ml inkwell bottles, after which 8 ml of culture was pelleted by centrifugation at room temperature for 5 min at 2,600 × *g*. The cell pellet was resuspended in 1 ml ice-cold extraction solvent containing 20:40:40 (v/v) water/acetonitrile/methanol. This was then transferred to approximately 500 μl of 0.1 mm zirconia/silica beads (BioSpec) and lysed by bead beating as above. Samples were centrifuged, and the extracts filtered through a 0.22 μm Nylon Spin-X microcentrifuge filter (Corning) and stored at -80 °C until analysis. ATP quantification was performed using an Agilent 1290 Infinity II liquid chromatography system coupled with an Agilent 6545 quadrupole time-of-flight (Q-TOF) mass spectrometer equipped with a Dual Agilent Jet Stream Electrospray Ionization (Dual AJS ESI) source operated in negative mode. Metabolites were separated on an InfinityLab Poroshell 120 HILIC-Z, 2.1 x 150 mm, 2.7 µm, 100 Å column (Agilent) as previously described.^25,56^ Mass spectra were recorded in profile mode from *m*/*z* 60 to 1200 using an acquisition rate of 1 spectra/sec in the 2 GHz extended dynamic range mode and 1700 *m*/*z* low mass range, using the sensitive slicer mode. The ESI and Q-TOF settings were optimized for the detection of ATP and were set as follows: gas temperature 350 °C, gas flow 13 l/min, sheath gas temperature 350 °C, sheath gas flow 12 l/min, and the capillary, nozzle, fragmentor, skimmer, and octopole voltages were 3500, 2000, 150, 45, and 750 V, respectively. Dynamic mass axis calibration was achieved by continuous infusion of a reference mass solution using an isocratic pump with a 100:1 splitter. HPLC-grade water (Cen-Med Enterprises) and LC-MS grade solvents (Fisher Chemical) were used for both the LC-MS mobile phase and metabolite extraction. Data analysis was performed using the Agilent MassHunter Qualitative (v10.0) and Quantitative Analysis Software (v10.1). ATP identification was based on mass retention times determined using a chemical standard (Sigma) and isotope distribution patterns, and quantified by integrating the area under the curve using a mass tolerance of 20 ppm. Calibration curves of ATP standards in extraction buffer and spiked into a homologous mycobacterial extract were used to determine the linear range and check for ion suppression effects.

### Drug resistance and viability assays

Resistance of *Mtb* strains to LOPAC hits and other compounds was assessed using the microbroth dilution method to determine MICs. Assays were prepared in flat-bottom 96-well microtire plates by either performing 2-fold serial dilutions in media at 2 × final drug concentration, or by spotting 2 μl of a 100 × drug dilution in DMSO followed by the addition of 98 μl of media. Logarithmic-phase subcultures were diluted to OD_600_ 0.01, and 100 μl was then added to the plate to give a final OD_600_ of 0.005 and 1 × drug concentration. Only the inner wells were used for assay, and the outer wells were aliquoted with 200 μl PBS or media. Checkerboard assays were prepared by spotting 2 μl of a 100 × dilution series of either linezolid or THL in DMSO down the plate columns and a 100 × dilution series of isoniazid in water across the rows, creating a two-dimensional matrix. Media and cells were then added as for MIC assays. Plates were incubated at 37 °C with gentle shaking (80–100 rpm), and bacterial growth was measured after 10 days by OD_600_ using a microplate reader. MIC data were normalized to drug-free control wells and fit with non-linear regression in Prism (v10.6.1) (GraphPad Software). The MIC was defined as the concentration in the well that resulted in 90% inhibition of bacterial growth. The fractional inhibitory concentration index (FICI) for checkboard assays was calculated as follows: FICI = (MIC_AB_/MIC_A_) + (MIC_BA_/MIC_B_).^57^ FICI values were calculated using wells closest to half the MIC for each drug, and the mean value reported.

Bacterial viability before and after drug exposure was determined by plating 100 µl of 1:10 serial dilutions in PBS with 0.05% tyloxapol onto agar. Colonies were counted after 3–4 weeks of incubation at 37 ºC.

### Statistical analysis

Statistical analyses were performed using Prism (v10.6.1) (GraphPad Software). Significance was determined by unpaired *t*-tests or ordinary one- or two-way ANOVA with testing for multiple comparisons as specified in the figure legends. CFU and ATP data were log_10_ transformed as indicated to improve normality and homogeneity of variance. The Brown-Forsythe and Kolmogorov-Smirnov tests, as well as the homoscedasticity and QQ plots, were used to assess the effects of data transformation and test assumptions.

## Acknowledgments

We acknowledge support from the following grants: NIH R01AI175972, R01AI139465 and R01AI173328 for C.V.M., J.C., and M.B., and R21AI159348 for A.C. and M.B., as well as T32AI070117 for A.C.

C.V.M., K.P., and M.B. conceived and designed the study. C.V.M., B.S.L., A.C., and J.C. performed the experiments. C.V.M. and B.S.L. analyzed the data. M.B. and K.P. provided resources. C.V.M. and M.B. wrote the paper. B.S.L. and K.P. critically reviewed and edited the paper.

## References

1. WHO (2024). Global tuberculosis report 2024. Geneva: World Health Organization, Liscence: CC BY-NC-SA 3.0 IGO.

2. Conyers, L.E., and Saunders, B.M. (2024). Treatment for non-tuberculous mycobacteria: challenges and prospects. Front Microbiol 15, 1394220. 10.3389/fmicb.2024.1394220.

3. Zeng, S., Soetaert, K., Ravon, F., Vandeput, M., Bald, D., Kauffmann, J.M., Mathys, V., Wattiez, R., and Fontaine, V. (2019). Isoniazid Bactericidal Activity Involves Electron Transport Chain Perturbation. Antimicrob Agents Chemother 63. 10.1128/AAC.01841-18.

4. Shetty, A., and Dick, T. (2018). Mycobacterial Cell Wall Synthesis Inhibitors Cause Lethal ATP Burst. Front Microbiol 9, 1898. 10.3389/fmicb.2018.01898.

5. Koul, A., Vranckx, L., Dhar, N., Gohlmann, H.W., Ozdemir, E., Neefs, J.M., Schulz, M., Lu, P., Mortz, E., McKinney, J.D., et al. (2014). Delayed bactericidal response of Mycobacterium tuberculosis to bedaquiline involves remodelling of bacterial metabolism. Nat Commun 5, 3369. 10.1038/ncomms4369.

6. Rao, S.P., Alonso, S., Rand, L., Dick, T., and Pethe, K. (2008). The protonmotive force is required for maintaining ATP homeostasis and viability of hypoxic, nonreplicating Mycobacterium tuberculosis. Proc Natl Acad Sci U S A 105, 11945–11950. 10.1073/pnas.0711697105.

7. Xu, Y., Ehrt, S., Schnappinger, D., and Beites, T. (2023). Synthetic lethality of Mycobacterium tuberculosis NADH dehydrogenases is due to impaired NADH oxidation. mBio 14, e0104523. 10.1128/mbio.01045-23.

8. Pethe, K., Sequeira, P.C., Agarwalla, S., Rhee, K., Kuhen, K., Phong, W.Y., Patel, V., Beer, D., Walker, J.R., Duraiswamy, J., et al. (2010). A chemical genetic screen in Mycobacterium tuberculosis identifies carbon-source-dependent growth inhibitors devoid of in vivo efficacy. Nat Commun 1, 57. 10.1038/ncomms1060.

9. Rebollo-Lopez, M.J., Lelievre, J., Alvarez-Gomez, D., Castro-Pichel, J., Martinez-Jimenez, F., Papadatos, G., Kumar, V., Colmenarejo, G., Mugumbate, G., Hurle, M., et al. (2015). Release of 50 new, drug-like compounds and their computational target predictions for open source anti-tubercular drug discovery. PLoS One 10, e0142293. 10.1371/journal.pone.0142293.

10. Mak, P.A., Rao, S.P., Ping Tan, M., Lin, X., Chyba, J., Tay, J., Ng, S.H., Tan, B.H., Cherian, J., Duraiswamy, J., et al. (2012). A high-throughput screen to identify inhibitors of ATP homeostasis in non-replicating Mycobacterium tuberculosis. ACS Chem Biol 7, 1190–1197. 10.1021/cb2004884.

11. March, V.F.A., Maghradze, N., McHedlishvili, K., Avaliani, T., Aspindzelashvili, R., Avaliani, Z., Kipiani, M., Tukvadze, N., Jugheli, L., Bouaouina, S., et al. (2025). Drug-induced differential culturability in diverse strains of Mycobacterium tuberculosis. Sci Rep 15, 3588. 10.1038/s41598-024-85092-7.

12. Zeng, S., Yang, D., Rens, C., and Fontaine, V. (2023). The Mycobacterium bovis BCG GroEL1 Contributes to Isoniazid Tolerance in a Dormant-Like State Model. Microorganisms 11. 10.3390/microorganisms11020286.

13. Vargas-Blanco, D.A., Zhou, Y., Zamalloa, L.G., Antonelli, T., and Shell, S.S. (2019). mRNA Degradation Rates Are Coupled to Metabolic Status in Mycobacterium smegmatis. mBio 10. 10.1128/mBio.00957-19.

14. Lee, B.S., Kalia, N.P., Jin, X.E.F., Hasenoehrl, E.J., Berney, M., and Pethe, K. (2019). Inhibitors of energy metabolism interfere with antibiotic-induced death in mycobacteria. J Biol Chem 294, 1936–1943. 10.1074/jbc.RA118.005732.

15. Ames, L., Allen, R., Boshoff, H.I.M., Cleghorn, L.A.T., Engelhart, C.A., Schnappinger, D., and Parish, T. (2025). Common Biological Properties of Mycobacterium tuberculosis MmpL3 Inhibitors. ACS Infect Dis 11, 2523–2533. 10.1021/acsinfecdis.5c00394.

16. Lindman, M., and Dick, T. (2019). Bedaquiline Eliminates Bactericidal Activity of beta-Lactams against Mycobacterium abscessus. Antimicrob Agents Chemother 63. 10.1128/AAC.00827-19.

17. Vilchèze, C. (2020). Mycobacterial Cell Wall: A Source of Successful Targets for Old and New Drugs. Applied Sciences 10. 10.3390/app10072278.

18. Nguyen, M.H., Haas, M.K., Kasperbauer, S.H., Calado Nogueira de Moura, V., Eddy, J.J., Mitchell, J.D., Khare, R., Griffith, D.E., Chan, E.D., and Daley, C.L. (2024). Nontuberculous Mycobacterial Pulmonary Disease: Patients, Principles, and Prospects. Clin Infect Dis 79, e27–e47. 10.1093/cid/ciae421.

19. Saukkonen, J.J., Duarte, R., Munsiff, S.S., Winston, C.A., Mammen, M.J., Abubakar, I., Acuña-Villaorduña, C., Barry, P.M., Bastos, M.L., Carr, W., et al. (2025). Updates on the Treatment of Drug-Susceptible and Drug-Resistant Tuberculosis: An Official ATS/CDC/ERS/IDSA Clinical Practice Guideline. American Journal of Respiratory and Critical Care Medicine 211, 15–33. 10.1164/rccm.202410-2096ST

20. McNeil, M.B., Chettiar, S., Awasthi, D., and Parish, T. (2019). Cell wall inhibitors increase the accumulation of rifampicin in Mycobacterium tuberculosis. Access Microbiol 1, e000006. 10.1099/acmi.0.000006.

21. Soetaert, K., Rens, C., Wang, X.M., De Bruyn, J., Laneelle, M.A., Laval, F., Lemassu, A., Daffe, M., Bifani, P., Fontaine, V., and Lefevre, P. (2015). Increased Vancomycin Susceptibility in Mycobacteria: a New Approach To Identify Synergistic Activity against Multidrug-Resistant Mycobacteria. Antimicrob Agents Chemother 59, 5057–5060. 10.1128/AAC.04856-14.

22. Rens, C., Laval, F., Daffe, M., Denis, O., Frita, R., Baulard, A., Wattiez, R., Lefevre, P., and Fontaine, V. (2016). Effects of Lipid-Lowering Drugs on Vancomycin Susceptibility of Mycobacteria. Antimicrob Agents Chemother 60, 6193–6199. 10.1128/AAC.00872-16.

23. Koh, E.I., Oluoch, P.O., Ruecker, N., Proulx, M.K., Soni, V., Murphy, K.C., Papavinasasundaram, K., Reames, C.J., Trujillo, C., Zaveri, A., et al. (2022). Chemical-genetic interaction mapping links carbon metabolism and cell wall structure to tuberculosis drug efficacy. Proc Natl Acad Sci U S A 119, e2201632119. 10.1073/pnas.2201632119.

24. Xu, W., DeJesus, M.A., Rucker, N., Engelhart, C.A., Wright, M.G., Healy, C., Lin, K., Wang, R., Park, S.W., Ioerger, T.R., et al. (2017). Chemical Genetic Interaction Profiling Reveals Determinants of Intrinsic Antibiotic Resistance in Mycobacterium tuberculosis. Antimicrob Agents Chemother 61. 10.1128/AAC.01334-17.

25. Mulholland, C.V., Wiggins, T.J., Cui, J., Vilcheze, C., Rajagopalan, S., Shultis, M.W., Reyes-Fernandez, E.Z., Jacobs, W.R., Jr., and Berney, M. (2024). Propionate prevents loss of the PDIM virulence lipid in Mycobacterium tuberculosis. Nat Microbiol 9, 1607–1618. 10.1038/s41564-024-01697-8.

26. Hermans, N., de Zwaan, R., Mulder, A., van den Dool, J., van Soolingen, D., Kremer, K., and Anthony, R. (2024). Mycobacterium tuberculosis complex sample processing by mechanical lysis, an essential step for reliable whole genome sequencing. J Microbiol Methods 227, 107053. 10.1016/j.mimet.2024.107053.

27. Kaser, M., Ruf, M.T., Hauser, J., Marsollier, L., and Pluschke, G. (2009). Optimized method for preparation of DNA from pathogenic and environmental mycobacteria. Appl Environ Microbiol 75, 414–418. 10.1128/AEM.01358-08.

28. Kapoor, R., and Yadav, J.S. (2010). Development of a rapid ATP bioluminescence assay for biocidal susceptibility testing of rapidly growing mycobacteria. J Clin Microbiol 48, 3725–3728. 10.1128/JCM.01482-10.

29. Sambandamurthy, V.K., Derrick, S.C., Hsu, T., Chen, B., Larsen, M.H., Jalapathy, K.V., Chen, M., Kim, J., Porcelli, S.A., Chan, J., et al. (2006). Mycobacterium tuberculosis DeltaRD1 DeltapanCD: a safe and limited replicating mutant strain that protects immunocompetent and immunocompromised mice against experimental tuberculosis. Vaccine 24, 6309–6320. 10.1016/j.vaccine.2006.05.097.

30. Mukherjee, D., Wu, M.L., Teo, J.W.P., and Dick, T. (2017). Vancomycin and Clarithromycin Show Synergy against Mycobacterium abscessus In Vitro. Antimicrob Agents Chemother 61. 10.1128/AAC.01298-17.

31. Peteroy, M., Severin, A., Zhao, F., Rosner, D., Lopatin, U., Scherman, H., Belanger, A., Harvey, B., Hatfull, G.F., Brennan, P.J., and Connell, N.D. (2000). Characterization of a Mycobacterium smegmatis mutant that is simultaneously resistant to D-cycloserine and vancomycin. Antimicrob Agents Chemother 44, 1701–1704. 10.1128/AAC.44.6.1701-1704.2000.

32. Neville, L.F., Shalit, I., Warn, P.A., Scheetz, M.H., Sun, J., Chosy, M.B., Wender, P.A., Cegelski, L., and Rendell, J.T. (2021). In Vivo Targeting of Escherichia coli with Vancomycin-Arginine. Antimicrob Agents Chemother 65. 10.1128/AAC.02416-20.

33. Dulberger, C.L., Rubin, E.J., and Boutte, C.C. (2020). The mycobacterial cell envelope - a moving target. Nat Rev Microbiol 18, 47–59. 10.1038/s41579-019-0273-7.

34. Goins, C.M., Sudasinghe, T.D., Liu, X., Wang, Y., O’Doherty, G.A., and Ronning, D.R. (2018). Characterization of Tetrahydrolipstatin and Stereoderivatives on the Inhibition of Essential Mycobacterium tuberculosis Lipid Esterases. Biochemistry 57, 2383–2393. 10.1021/acs.biochem.8b00152.

35. Ravindran, M.S., Rao, S.P., Cheng, X., Shukla, A., Cazenave-Gassiot, A., Yao, S.Q., and Wenk, M.R. (2014). Targeting lipid esterases in mycobacteria grown under different physiological conditions using activity-based profiling with tetrahydrolipstatin (THL). Mol Cell Proteomics 13, 435–448. 10.1074/mcp.M113.029942.

36. Belardinelli, J.M., Larrouy-Maumus, G., Jones, V., Sorio de Carvalho, L.P., McNeil, M.R., and Jackson, M. (2014). Biosynthesis and translocation of unsulfated acyltrehaloses in Mycobacterium tuberculosis. J Biol Chem 289, 27952–27965. 10.1074/jbc.M114.581199.

37. Kidwai, S., Bouzeyen, R., Chakraborti, S., Khare, N., Das, S., Priya Gosain, T., Behura, A., Meena, C.L., Dhiman, R., Essafi, M., et al. (2019). NU-6027 Inhibits Growth of Mycobacterium tuberculosis by Targeting Protein Kinase D and Protein Kinase G. Antimicrob Agents Chemother 63. 10.1128/AAC.00996-19.

38. Paik, S.R., Jault, J.M., and Allison, W.S. (1994). Inhibition and inactivation of the F1 adenosinetriphosphatase from Bacillus PS3 by dequalinium and activation of the enzyme by lauryl dimethylamine oxide. Biochemistry 33, 126–133. 10.1021/bi00167a016.

39. Mendling, W., Weissenbacher, E.R., Gerber, S., Prasauskas, V., and Grob, P. (2016). Use of locally delivered dequalinium chloride in the treatment of vaginal infections: a review. Arch Gynecol Obstet 293, 469–484. 10.1007/s00404-015-3914-8.

40. Hasenoehrl, E.J., Wiggins, T.J., and Berney, M. (2020). Bioenergetic Inhibitors: Antibiotic Efficacy and Mechanisms of Action in Mycobacterium tuberculosis. Front Cell Infect Microbiol 10, 611683. 10.3389/fcimb.2020.611683.

41. Russell, J.B., and Cook, G.M. (1995). Energetics of bacterial growth: balance of anabolic and catabolic reactions. Microbiol Rev 59, 48–62. 10.1128/mr.59.1.48-62.1995.

42. Vilcheze, C., and Jacobs, W.R., Jr. (2019). The Isoniazid Paradigm of Killing, Resistance, and Persistence in Mycobacterium tuberculosis. J Mol Biol 431, 3450–3461. 10.1016/j.jmb.2019.02.016.

43. De la Fuente, I.M., Cortes, J.M., Valero, E., Desroches, M., Rodrigues, S., Malaina, I., and Martinez, L. (2014). On the dynamics of the adenylate energy system: homeorhesis vs homeostasis. PLoS One 9, e108676. 10.1371/journal.pone.0108676.

44. Lamprecht, D.A., Finin, P.M., Rahman, M.A., Cumming, B.M., Russell, S.L., Jonnala, S.R., Adamson, J.H., and Steyn, A.J. (2016). Turning the respiratory flexibility of Mycobacterium tuberculosis against itself. Nat Commun 7, 12393. 10.1038/ncomms12393.

45. Liang, L., Lin, D., Chen, Y., Li, J., Liang, W., Zhao, H., Luo, W., Tian, G.B., and Feng, S. (2022). Single-Fluorescence ATP Sensor Based on Fluorescence Resonance Energy Transfer Reveals Role of Antibiotic-Induced ATP Perturbation in Mycobacterial Killing. mSystems 7, e0020922. 10.1128/msystems.00209-22.

46. Maglica, Z., Ozdemir, E., and McKinney, J.D. (2015). Single-cell tracking reveals antibiotic-induced changes in mycobacterial energy metabolism. mBio 6, e02236–02214. 10.1128/mBio.02236-14.

47. Quigley, J., and Lewis, K. (2022). Noise in a Metabolic Pathway Leads to Persister Formation in Mycobacterium tuberculosis. Microbiol Spectr 10, e0294822. 10.1128/spectrum.02948-22.

48. Akela, A.K., and Kumar, A. (2021). Bioenergetic Heterogeneity in Mycobacterium tuberculosis Residing in Different Subcellular Niches. mBio 12, e0108821. 10.1128/mBio.01088-21.

49. Lodhiya, T., Palande, A., Veeram, A., Larrouy-Maumus, G., Beste, D.J., and Mukherjee, R. (2025). ATP burst is the dominant driver of antibiotic lethality in Mycobacterium smegmatis. eLife 13. 10.7554/eLife.99656.3.

50. Wivagg, C.N., Bhattacharyya, R.P., and Hung, D.T. (2014). Mechanisms of beta-lactam killing and resistance in the context of Mycobacterium tuberculosis. J Antibiot (Tokyo) 67, 645–654. 10.1038/ja.2014.94.

51. Longo, B.M., Trunfio, M., and Calcagno, A. (2024). Dual beta-lactams for the treatment of Mycobacterium abscessus: a review of the evidence and a call to act against an antibiotic nightmare. J Antimicrob Chemother 79, 2731–2741. 10.1093/jac/dkae288.

52. Hugonnet, J.E., Tremblay, L.W., Boshoff, H.I., Barry, C.E., 3rd, and Blanchard, J.S. (2009). Meropenem-clavulanate is effective against extensively drug-resistant Mycobacterium tuberculosis. Science 323, 1215–1218. 10.1126/science.1167498.

53. Negatu, D.A., Aragaw, W.W., Dartois, V., and Dick, T. (2024). A pairwise approach to revitalize beta-lactams for the treatment of TB. Antimicrob Agents Chemother 68, e0003424. 10.1128/aac.00034-24.

54. Story-Roller, E., Maggioncalda, E.C., and Lamichhane, G. (2019). Select beta-Lactam Combinations Exhibit Synergy against Mycobacterium abscessus In Vitro. Antimicrob Agents Chemother 63. 10.1128/AAC.02613-18.

55. Snapper, S.B., Melton, R.E., Mustafa, S., Kieser, T., and Jacobs, W.R., Jr. (1990). Isolation and characterization of efficient plasmid transformation mutants of Mycobacterium smegmatis. Mol Microbiol 4, 1911–1919.

56. Dai, Y., and Hsiao, J.J. (2019). Discovery Metabolomics LC/MS Methods Optimized for Polar Metabolites. Application note, Agilent Technologies, Inc.

57. Hsieh, M.H., Yu, C.M., Yu, V.L., and Chow, J.W. (1993). Synergy assessed by checkerboard. A critical analysis. Diagn Microbiol Infect Dis 16, 343–349. 10.1016/0732-8893(93)90087-n.

