## Supplementary Information for "The mycobacterial ATP burst is a lysis artifact and serves as an assay for drug-induced cell wall damage"

### Table of contents

#### Supplementary Figures

S1. The isoniazid-induced ATP burst is not observed in heat-inactivated cultures.
S2. The ATP burst is an experimental artifact.
S3. Quantification of ATP by LC-MS.
S4. Optimization of ATP screening for identification of cell wall inhibitors.
S5. Cross-resistance of *Mtb* to isoniazid and nialamide.
S6. Effect of LOPAC hits on the growth of virulent *Mtb* H37Rv.
S7. Effect of co-treatment of *Mtb* with isoniazid and bioenergetics inhibitors on ATP and survival.
S8. *Mtb* growth optimization in 96-well plate format.
S9. PDIM-positive *Mtb* shows an artifactual ATP burst with isoniazid.
S10. Whole-cell ATP dose-response of PDIM-positive *Mtb* to tetrahydrolipstatin and vancomycin.

#### Supplementary Tables

S1. $MIC_{90}$ values and concentrations of compounds used in the isoniazid co-treatment assay.
S2. FICI assays for tetrahydrolipstatin and linezolid with isoniazid.
S3. Culture media reagents and supplements used in this study.
S4. Inhibitors used in this study.

#### Supplementary Figures

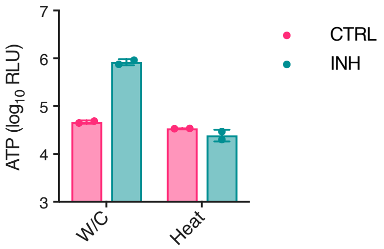


**Figure S1. The isoniazid-induced ATP burst is not observed in heat-inactivated cultures.** In a preliminary experiment, *Mtb* $mc^2$6230 at $OD_{600}$ 0.33 was treated with isoniazid (INH, 5 µg/ml) for 24 h, and ATP was measured by the BacTiter-Glo™ assay using either whole-cell cultures (W/C) or cultures inactivated by boiling at 100 °C for 30 minutes (Heat). $n = 2$ technical replicates.

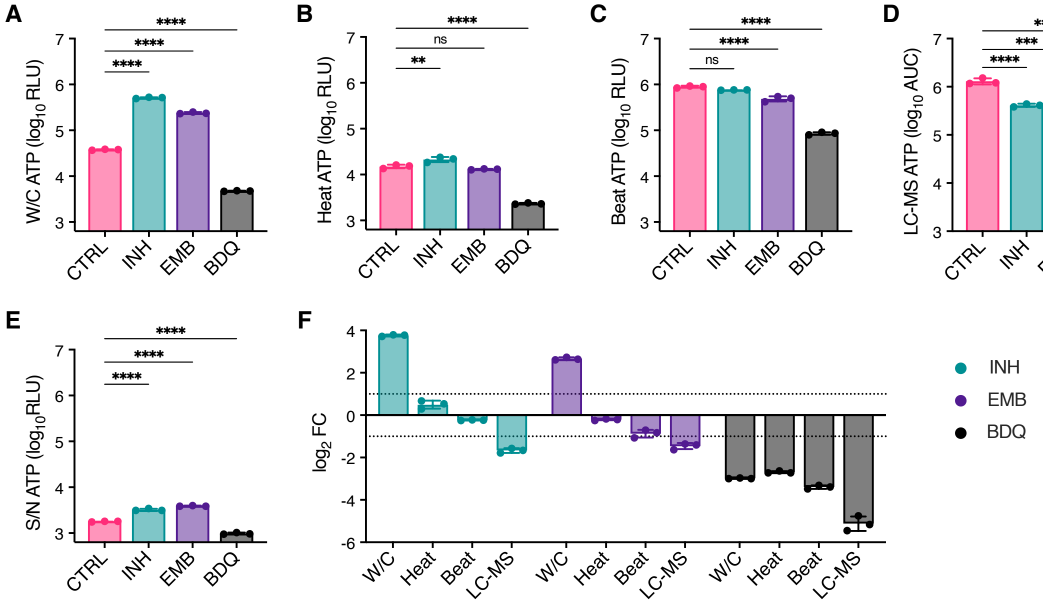


**Figure S2. The ATP burst is an experimental artifact.** *Mtb* mc[2]6230 at $OD_{600}$ 0.33 was treated with isoniazid (INH, 3 µg/ml), ethambutol (EMB, 50 µg/ml), and bedaquiline (BDQ, 12.5 µg/ml) for 24 h. **A–C**. ATP was measured using the BacTiter-Glo™ assay from either (A) whole-cell cultures ('W/C'), (B) the supernatant from cells lysed by heat inactivation ('Heat'), or (C) supernatant from cells lysed by bead beating ('Beat'). **D**. ATP levels in the same cultures measured by LC-MS. **E**. ATP in the culture supernatant measured using the BacTiter-Glo™ assay. **F**. Fold-change in ATP levels compared to untreated controls for the different methods shown in (A–D). The dotted lines indicate a 2-fold change. Mean ± SD, $n = 3$ replicate cultures. $^{**}P < 0.01$, $^{***}P < 0.001$, $^{****}P < 0.0001$; one-way ANOVA with Šídák's multiple comparison test (MCT). These data represent an independent repeat of the experiment shown in Fig. 1.

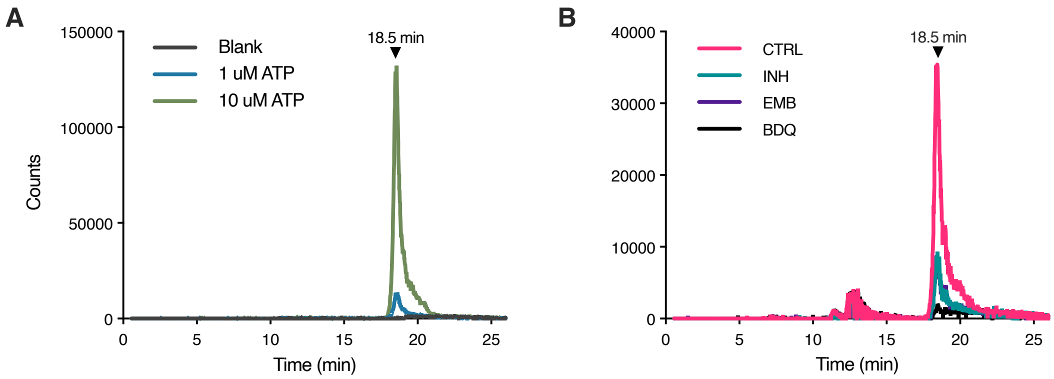


**Figure S3. Quantification of ATP by LC-MS. A–B.** Extracted ion chromatograms (EICs) for ATP, *m*/*z* = 505.9885, for (A) ATP compound standards and (B) sample extracts. One representative extract from each condition is shown.

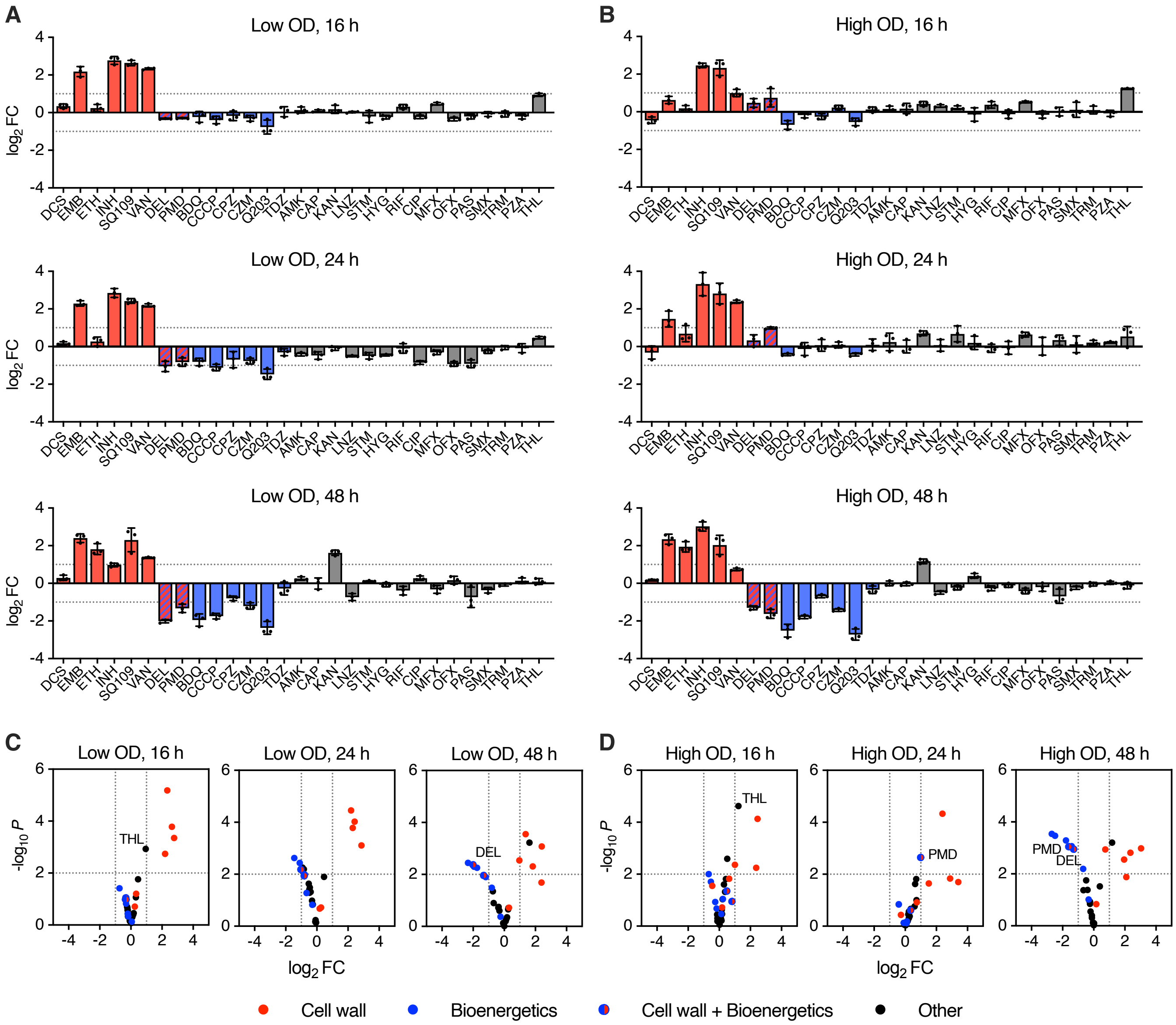


**Figure S4. Optimization of ATP screening for identification of cell wall inhibitors. A–B.** *Mtb* mc[2]6230 was diluted to (A) $OD_{600}$ 0.033 (low OD) or (B) $OD_{600}$ 0.33 (high OD) and treated with 28 reference compounds (see Table 1) at 10 µM alongside DMSO-only controls. Cultures were incubated in 96-well microtiter plates for 16, 24, or 48 h, and ATP was measured using the BacTiter-Glo™ whole-cell assay. $Log_2$ fold-changes relative to DMSO-only controls are shown. **C–D**. Volcano plots showing responses relative to DMSO-only controls for (C) the low-density cultures from (A) and (D) the high-density cultures from (B). Dotted lines indicate a 2-fold change and *P*-value of 0.01. Mean ± SD, *n* = 3 replicate wells. See also Fig. 3A–B, which represents an independent repeat of the low OD, 24 h experiment.

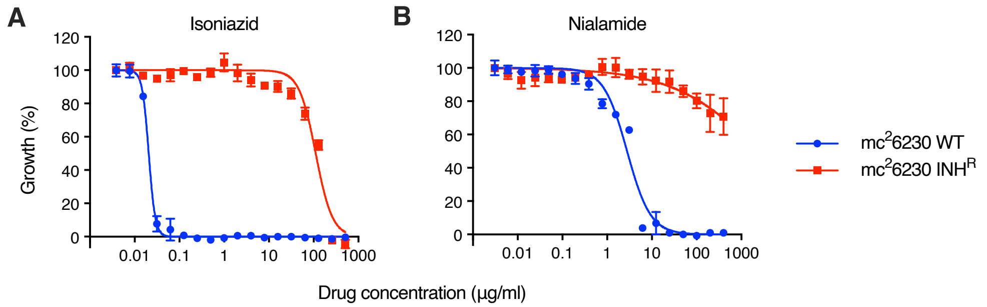


**Figure S5. Cross-resistance of *Mtb* to isoniazid and nialamide. A–B.** Minimum inhibitory concentration (MIC) assays of *Mtb* mc$^2$6230 wildtype (WT) and an isoniazid-resistant mutant (INH$^R$) with (A) isoniazid and (B) nialamide. Growth was measured by $OD_{600}$ and normalized to no-drug control wells. The isoniazid-resistant mutant has a *katG* 1627AG>A frameshift mutation, rendering it unable to activate isoniazid. Mean ± SD, $n = 2$ replicate wells.

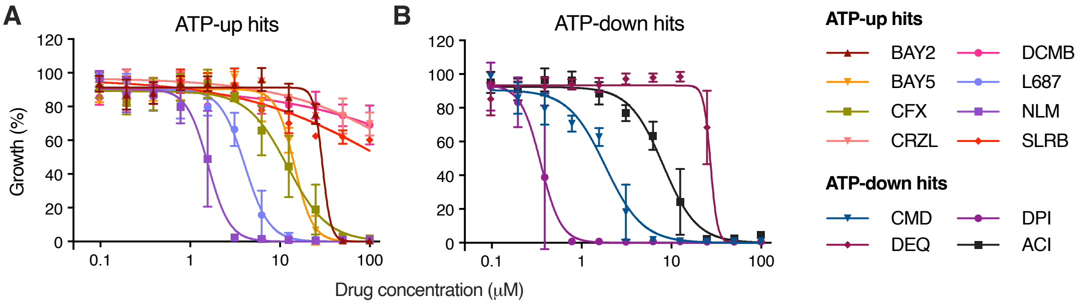


**Figure S6. Effect of LOPAC hits on the growth of virulent *Mtb* H37Rv. A–B.** MIC assays for PDIM-positive *Mtb* H37Rv AE1001 with selected LOPAC compounds that either (A) significantly increased the ATP signal by >2-fold ("ATP-up" hits) or (B) significantly decreased the ATP signal by > 80% ("ATP-down" hits) (see Table 2). The culture media was additionally supplemented with 0.1 mM propionate to support PDIM production. Growth was measured by $OD_{600}$ and normalized to no-drug control wells. Mean ± SD, *n* = 4 replicate wells from two independent experiments.

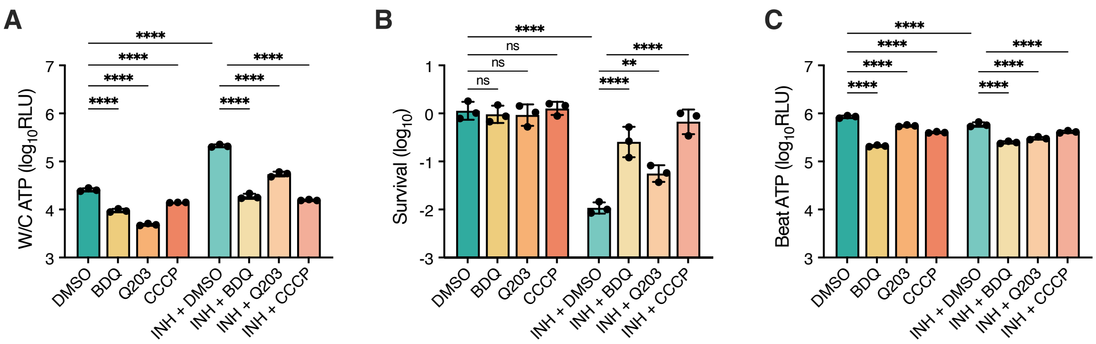


**Figure S7. Effect of co-treatment of *Mtb* with isoniazid and bioenergetics inhibitors on ATP and survival.** *Mtb* mc[2]6230 at $OD_{600}$ 0.33 was treated with isoniazid (INH, 0.1 μg/ml), bedaquiline (BDQ, 2.5 μg/ml), Q203 (200 nM), and CCCP (20 μg/ml) for 24 h. **A.** ATP measured using the BacTiter-Glo™ whole-cell assay. **B.** Relative survival following drug treatment. Cultures were plated for CFU before and after treatment to assess viability. The starting cell density was $2 \times 10^8$ CFU/ml. **C.** ATP measured using the BacTiter-Glo™ beat assay. Mean ± SD, $n = 3$ replicate cultures. $^{**}P < 0.01$, $^{****}P < 0.0001$; one-way ANOVA with Šídák's MCT. This experiment was repeated with similar results.

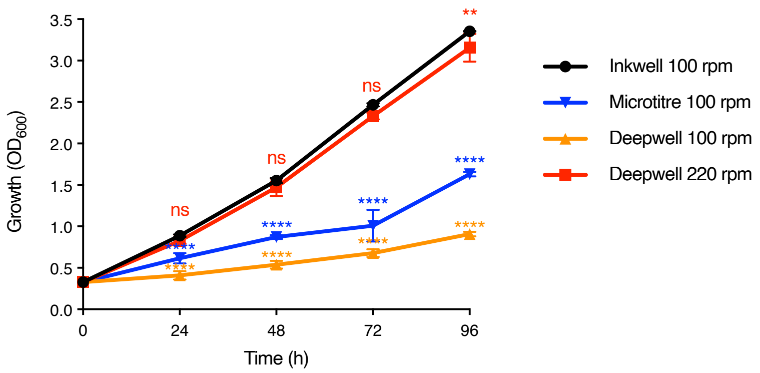


**Figure S8. *Mtb* growth optimization in 96-well plate format.** *Mtb* mc$^2$6230 cultures at OD 0.33 were grown in either inkwell bottles, a standard 96-well microtiter plate, or 2 ml V-bottom 96-deepwell blocks, shaken gently at 100 rpm or vigorously at 220 rpm. The culture volume was 5 ml for inkwells and 200 µl for 96-well plates and deepwell blocks. Deepwell blocks were sealed with sterile foil seals, contained in sealed bags and secured using the Khuner Duetz System (Basel, Switzerland) for fast shaking. Mean ± SD, $n = 3$ replicate inkwell cultures, and $n = 3–6$ replicate wells for 96-well plates and deepwell blocks. $^{**}P < 0.01$; $^{****}P < 0.0001$ vs. inkwell cultures; two-way ANOVA test with Dunnett's MCT. Deepwell blocks shaken at 220 rpm reproducibly resulted in growth comparable to inkwells.

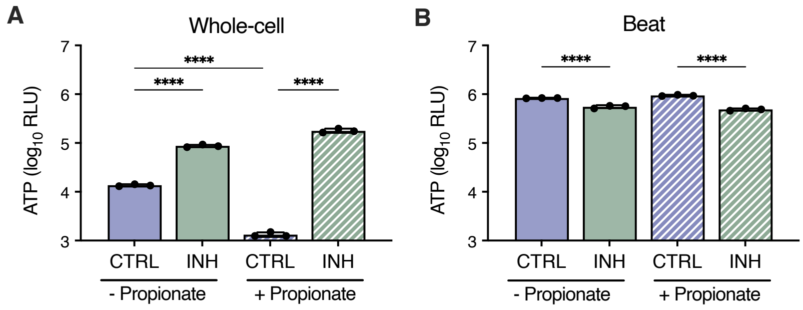


**Figure S9. PDIM-positive *Mtb* shows an artifactual ATP burst with isoniazid.** PDIM-positive *Mtb* mc$^2$6230 AE1601 was diluted to $OD_{600}$ of 0.05 into 7H9/OADC/glycerol/tyloxapol media with and without 0.1 mM propionate and treated with isoniazid for 24 h. **A–B.** ATP was measured using (A) the BacTiter-Glo™ whole-cell and (B) beat assays. Mean ± SD, $n = 3$ replicate cultures. ****$P < 0.0001$; one-way ANOVA with Šídák's MCT.

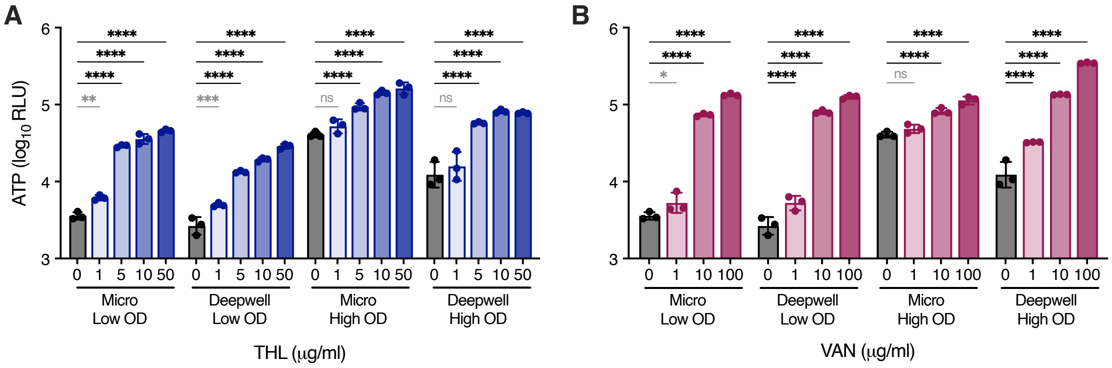


**Figure S10. Whole-cell ATP dose-response of PDIM-positive *Mtb* to tetrahydrolipstatin and vancomycin. A–B.** *Mtb* mc$^2$6230 AE1601 was diluted to either OD 0.033 (low OD) or OD 0.33 (high OD) and treated with different concentrations of (A) tetrahydrolipstatin or (B) vancomycin for 24 h, and ATP was measured using the BacTiter-Glo™ whole-cell assay. Cultures were treated in either standard 96-well microtiter plates with gentle shaking at 100 rpm or in 96-deepwell blocks with vigorous shaking at 220 rpm (see Fig. S8). Mean ± SD, $n = 3$ replicate wells. *$P < 0.05$, **$P < 0.01$, ***$P < 0.001$, ****$P < 0.0001$; two-way ANOVA with Dunnett's MCT. Comparisons with ≤2-fold changes are indicated in grey and >2-fold in black. Based on these results, we opted to use low-density cultures with 1 µg/ml of each drug in 96-well microtiter plates for the cotreatment assay in Fig. 5E, as this induced minimal response compared to DMSO-only (≤2-fold), while strong responses were elicited at higher concentrations.

#### Supplementary Tables

**Table S1. $MIC_{90}$ values and concentrations of compounds used in the isoniazid co-treatment assay.** Concentrations are reported in µg/ml, except for Q203, which is in nM. MICs were measured in *Mtb* $mc^2$6230 using the microbroth dilution method.

| Abbr. | Name | $MIC_{90}$ | conc1 | conc2 |
|---|---|---|---|---|
| INH | Isoniazid | 0.05 | N/A | N/A |
| AMK | Amikacin | 0.5 | 2 | 5 |
| BDQ | Bedaquiline | 0.25 | 0.5 | 1.25 |
| CCCP | Carbonyl cyanide 3-chlorophenylhydrazone | 10 | 20 | 50 |
| CZM | Clofazimine | 0.5 | 0.5 | 1.25 |
| LNZ | Linezolid | 1 | 2 | 5 |
| MFX | Moxifloxacin | 0.1 | 0.2 | 0.5 |
| PAS | 4-Aminosalicylic acid | 0.1 | 0.2 | 0.5 |
| Q203 | Q203 | 20 (nM) | 40 (nM) | 100 (nM) |
| RIF | Rifampicin | 0.05 | 0.1 | 0.25 |
| STM | Streptomycin | 1 | 4 | 10 |
| THL | Tetrahydrolipstatin | 16 – 32 | 50 | 125 |

**Table S2. FICI assays for tetrahydrolipstatin and linezolid with isoniazid.** Data are shown as growth normalized to no-drug control wells. Yellow wells indicate the $MIC_{90}$ for each drug alone. Wells nearest half the MIC for each drug (orange wells) were used to calculate the FIC index as follows: FICI = $(MIC_{AB}/MIC_A) + (MIC_{BA}/MIC_B)$. Data show mean growth from two replicate plates.

| | | **INH** (μg/ml) | | | | | | | | | |
|---|---|---|---|---|---|---|---|---|---|---|---|
| | | **0.4** | **0.2** | **0.1** | **0.05** | **0.025** | **0.0125** | **0.0063** | **0.0031** | **0.0016** | **0** |
| **THL (μg/ml)** | **32** | 0.0% | -3.9% | 1.2% | -0.8% | 1.2% | 1.2% | 1.6% | 1.7% | 1.8% | 0.0% |
| | **16** | 1.0% | 0.3% | 1.5% | 0.7% | 1.4% | 2.4% | 3.0% | 3.6% | 3.4% | 2.6% |
| | **8** | 0.1% | 0.1% | -0.7% | 0.0% | 0.7% | 4.8% | 13.7% | 12.6% | 11.8% | 13.4% |
| | **4** | -1.2% | -0.1% | -0.7% | -2.7% | -0.3% | 13.6% | 25.8% | 36.3% | 37.3% | 40.2% |
| | **2** | -0.3% | -1.5% | -0.4% | -1.1% | -0.7% | 39.6% | 44.3% | 46.9% | 47.2% | 48.2% |
| | **0** | -1.3% | -2.0% | -0.3% | -1.1% | 60.2% | 92.1% | 94.6% | 98.5% | 102.0% | 97.8% |

| | | **INH** (μg/ml) | | | | | | | | | |
|---|---|---|---|---|---|---|---|---|---|---|---|
| | | **0.4** | **0.2** | **0.1** | **0.05** | **0.025** | **0.0125** | **0.0063** | **0.0031** | **0.0016** | **0** |
| **LNZ (μg/ml)** | **1** | 0.0% | -0.4% | 0.4% | 0.9% | 1.1% | 2.0% | 6.1% | 8.0% | 3.6% | 4.3% |
| | **0.5** | 0.1% | 0.3% | 2.0% | 1.3% | 1.9% | 54.7% | 66.9% | 72.8% | 52.4% | 67.8% |
| | **0.3** | -0.5% | 0.5% | -0.3% | 0.6% | 34.4% | 85.5% | 88.5% | 89.7% | 81.8% | 94.4% |
| | **0.1** | -0.5% | 0.1% | 1.0% | -0.8% | 70.8% | 101.4% | 99.7% | 100.7% | 96.9% | 107.8% |
| | **0.1** | -0.2% | -0.2% | 0.7% | 0.8% | 72.8% | 100.2% | 101.2% | 103.7% | 103.8% | 107.4% |
| | **0** | -1.1% | -1.4% | -0.6% | -0.7% | 67.6% | 92.6% | 102.2% | 103.8% | 102.2% | 106.3% |

**Table S3. Culture media reagents and supplements used in this study.**

| Name | Supplier | Catalog # |
|---|---|---|
| Bovine serum albumin fraction V | GoldBio | A-420-1 |
| Catalase | Sigma-Aldrich | C1345 |
| D-Calcium pantothenate | Acros Organics | 243305000 |
| Dextrose (D-glucose) | Fisher Chemical | D16 |
| Glycerol | Fisher Chemical | G33 |
| LB Agar | BD Difco | 244520 |
| LB Broth | BD Difco | 244610 |
| Middlebrook 7H10 | BD Difco | 262710 |
| Middlebrook 7H9 | BD Difco | 271310 |
| Middlebrook ADC Enrichment | BD Difco | 211887 |
| Sodium chloride | Fisher Chemical | S271 |
| Sodium oleate | Strem Chemicals | 11-1280 |
| Sodium propionate | Sigma-Aldrich | P1880 |
| Tween 80 | Sigma-Aldrich | P1754 |
| Tyloxapol | Sigma-Aldrich | T8761 |

**Table S4. Inhibitors used in this study.**

| Abbr. | Full compound name | Manufacturer | Catalog # |
|---|---|---|---|
| AMK | Amikacin disulfate salt | Acros Organics | 455190050 |
| BDQ | Bedaquiline | Gift from Kevin Pethe, Nanyang Technological University | |
| CAP | Capreomycin Sulfate | Alfa Aesar | J66684MD |
| CCCP | Carbonyl cyanide 3-chlorophenylhydrazone | Tocris | 0452 |
| CIP | Ciprofloxacin | Acros Organics | 449620050 |
| CPZ | Chlorpromazine hydrochloride | MP Biomedicals, Inc. | 190326 |
| CZM | Clofazimine | TCI | C2866 |
| DCS | D-Cycloserine | Acros Organics | 228480050 |
| DEL | Delamanid | MedChemExpress | HY-10846 |
| EMB | Ethambutol dihydrochloride | Alfa Aesar | J60695 |
| ETH | Ethionamide | TCI | E0695 |
| HYG | Hygromycin B | Invitrogen | 10687010 |
| INH | Isoniazid | Sigma | I-3377 |
| KAN | Kanamycin Sulfate | Fisher BioReagents | BP906 |
| LNZ | Linezolid | Acros Organics | 460592500 |
| MFX | Moxifloxacin hydrochloride | Acros Organics | 457960010 |
| OFX | Ofloxacin | Alfa Aesar | J62080 |
| PAS | 4-Aminosalicylic acid | Acros Organics | 104620050 |
| PMD | Pretomanid | MedChemExpress | HY-10844 |
| Q203 | Q203 | Gift from Kevin Pethe, Nanyang Technological University | |
| RIF | Rifampicin | Sigma | R3501 |
| SMX | Sulfamethoxazole | Fluka analytica-Sigma | S7507 |
| SQ109 | SQ109 | Sigma | SML1309 |
| STM | Streptomycin sulfate salt | Gold Biotechnology | S15050 |
| TDZ | Thioridazine hydrochloride | Tocris | 3070 |
| THL | Tetrahydrolipstatin (Orlistat) | TCI | O0381 |
| TRM | Trimethoprim | Teknova | T1205 |
| VAN | Vancomycin hydrochloride | Alfa Aesar | J62790 |
